# Novel genomic offset metrics account for local adaptation in climate suitability forecasts and inform assisted migration

**DOI:** 10.1101/2023.06.05.541958

**Authors:** Susanne Lachmuth, Thibaut Capblancq, Anoob Prakash, Stephen R. Keller, Matthew C. Fitzpatrick

## Abstract

Local adaptation is increasingly being integrated into macroecological models, offering an evolutionary perspective that has been largely missing from climate change biogeography. Genomic offsets, which quantify the disruption of existing genotype-environment associations under environmental change, are an informative landscape genomic tool that allows for the incorporation of intra-specific adaptive differentiation into forecasts of climate suitability and thus management planning. Gradient forest (GF), a method originally developed to model community turnover along environmental gradients, is now the most commonly used approach in genomic offset estimation. However, major hurdles in the application of GF-derived offsets are (1) an inability to interpret the absolute magnitude of genomic offsets in a biologically meaningful way and (2) uncertainty in how genomic offsets compare to established species-level approaches like Ecological Niche Models (ENMs).

We used both ENMs and novel, genomic offset metrics derived from GF modeling of genomic variation along climatic gradients to assess the climate change vulnerability of red spruce (*Picea rubens*), a cool-temperate tree species endemic to eastern North America. We show how genomic offsets can be standardized relative to contemporary genomic variation across the landscape to better represent their biological significance and facilitate comparisons among studies. In three common gardens, we found a significant negative relationship between standardized genomic offsets and red spruce growth and higher explanatory power for standardized offsets than (raw) climate transfer distances.

We also derived new threshold-based metrics that we term Donor and Recipient Importance and which quantify transferability of propagules between donor and recipient localities while minimizing disruption of genotype-environment associations. This approach leverages landscape genomic information to account for local adaptation when predicting climate suitability. ENMs and our novel genomic offset metrics largely agreed in forecasting drastic northward range shifts. Combining several offset-based metrics, we show that the projected northward shift of suitable climate mainly applies to populations located in the center and northern parts of the current range, whereas southern populations might be able to persist *in situ* owing to specific local climate adaptations. The novel metrics thus yield refined, region-specific prognoses for local persistence and show how management could be improved by considering assisted migration.

## Introduction

Anthropogenic climate change is disrupting local adaptation to climate at an unprecedented pace, with cascading effects on individual fitness, population demography (Etterson et al. 2020; Shaw and Etterson 2012), and ecosystem functioning and services (Schröter et al. 2005). For species like forest trees with long generation time and limited dispersal, the rate of climate change will almost certainly exceed the ability of populations to realize rapid *in situ* evolutionary adaptation or track favorable climates by dispersing to new locations (Aitken et al. 2008; Dauphin et al. 2021). Climate-conscious management may therefore need to move beyond *in situ* conservation and place a greater emphasis on assisted migration programs that encompass both assisted gene flow within and assisted colonization beyond the current range (definitions according to Aitken and Bemmels (2016), Aitken and Whitlock (2013)). These management actions would first require rethinking the ‘local is best’ paradigm, which assumes local sources will continue to maintain higher fitness compared to non-local sources due to current local adaptation. However, in rapidly changing environments, a “non-local is best” approach may be preferred and will require new concepts that leverage provenance trials and genomic resources to identify seed sources that minimize disruption of local adaptation to climate (Aitken and Bemmels 2016; Etterson et al. 2020; Isabel, Holliday, and Aitken 2020).

Ultimately, decisions on whether to implement artificial transfer of propagules and how to frame such transfers (e.g., within or beyond a species contemporary range) are complex and require careful consideration of risks and benefits (Hagerman, Kozak, and Dalrymple 2021). Combining ecological and evolutionary research can help to assess potential translocations and maximize their success. Most studies to date have relied on provenance trials and ecological niche models (ENMs) for matching seed sources with potential planting sites (Rellstab, Dauphin, and Exposito-Alonso 2021). Whereas provenance trials are costly and often restricted to few populations (Aitken and Bemmels 2016), the environmental and species data required for fitting ENMs often are widely available and the models have become one of the most common quantitative ecological methods not only in academia, but also for conservation practitioners (Pacifici et al. 2015; Zurell et al. 2020). However, ENMs often lack an evolutionary perspective, as they do not account for intra-specific genetic differentiation, including potentially important differences between populations in climate adaptation. ENM studies that have incorporated intra-specific variability report better predictive performance (Chardon et al. 2020) and often lower vulnerability to climate change than species-level ENMs (Benito Garzón et al. 2011; Bush et al. 2016; Razgour et al. 2019).

Recently, the concept of genomic offset (first developed by Fitzpatrick and Keller (2015), reviewed by Capblancq, Fitzpatrick, et al. (2020)) has emerged as a promising tool for incorporating genomic information into climate change impact assessments. Based on model predictions of genome-wide allele frequency variation along environmental gradients, genomic offset quantifies the expected disruption of existing genotype-environment associations (hereafter G - E disruption) under environmental change. To be most informative and to avoid confounding effects of non-adaptive genetic differentiation, genomic offsets ideally should be based on loci that have been identified as potentially adaptive *via* genotype-environment association (GEA) studies and should be validated by analyzing the fitness effects of the estimated G – E disruption, ideally in controlled experiments (Capblancq, Fitzpatrick, et al. 2020; Fitzpatrick et al. 2021; Rellstab, Dauphin, and Exposito-Alonso 2021). Further, genomic offsets have great untapped potential for advancing predictive provenancing in genomically-informed assisted migration. However, genomic offsets, as originally introduced by Fitzpatrick and Keller (2015), did not consider the possibility of migration, which is crucial for planning translocations. To assess the relative potentials of adaptation and migration in mitigating climate change risks, Gougherty, Keller, and Fitzpatrick (2021) defined three offset-based metrics: “Local”, “Forward” and “Reverse Offset”. Local Offset (LO) is the same metric described by Fitzpatrick and Keller (2015) and is defined as the G - E disruption a population would experience *in situ* under an anticipated level of environmental change and thus informs about the necessity of emigration or intervention through assisted gene migration. Forward Offset represents the residual G - E disruption a population would experience if it was transferred (or could migrate) to the location *most similar* to its current genomic adaptation under future environmental conditions within a pre-defined geographic area. Similarly, Reverse Offset quantifies the residual G - E disruption the population most-pre-adapted to the future environmental conditions at a given location would still experience after transfer or immigration. Collectively, these offset metrics allow visualization and evaluation of where G - E disruption is expected to be most severe, where populations are predicted to be poorly adapted to all projected environmental conditions in the study area, and where rescue through (assisted) migration is expected to be either most successful or constrained (Gougherty, Keller, and Fitzpatrick 2021).

For application in management planning, what is still needed is a fuller consideration of how existing populations differ in terms of their ability to contribute preadapted alleles to future populations (i.e., their differing potential to serve as donors) and the identification of locations for which genomic offsets remain within a biologically tolerable range (the potential of a location to benefit from receiving propagules from current populations, i.e., to serve as recipients). However, consideration of intra-specific differentiation in climate change response introduces a new dilemma – namely, the need to place the absolute magnitude of genomic offsets in a biologically relevant context. Because genomic offsets derived from Gradient Forest modeling (hereafter, GF offsets) are unitless distances, they lack direct biological interpretation. Although it is possible to compare offsets between populations within the same study and assess relative risk, it is not possible to determine *a priori* what level of GF offset will be tolerable for a population without a link to an expected biological response. To address this problem, we propose the standardization of GF offsets relative to the currently observed spatial genomic variation across a species’ contemporary range. In other words, we ask: where do the magnitudes of predicted GF offsets for a given change in environment (through time or space) fall within the distribution of GF offsets observed between contemporary populations across space? The standardized GF offsets provide a more objective means to set a threshold for ‘tolerable’ genomic offsets and provide an objective basis to estimate propagule transferability and to assess the importance of each population to serve as a donor (i.e., seed source) and of any geographic location to serve as recipient (i.e., potential planting location) of propagules (Figure 1).

**Figure 1:**
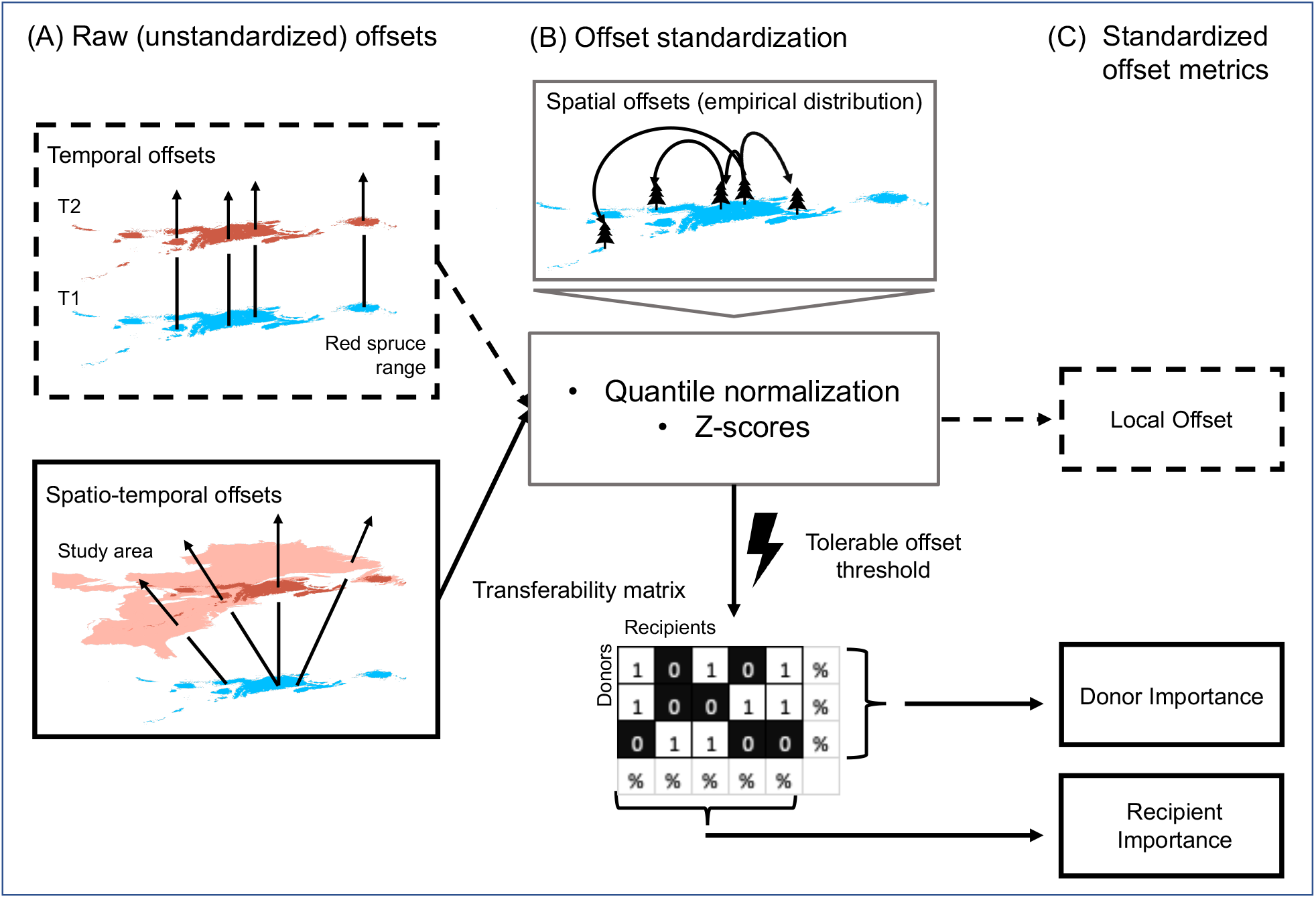
Schematic illustration of the calculation of standardized genomic offset metrics. (A) Standardization is applied to two types of genomic offsets to produce our standardized offset metrics: Temporal offsets quantify the *in situ* genomic offset expected under a change from current (blue) to future (red) climate in each grid cell in the current red spruce range; Spatio-temporal offsets describe the G - E disruption for a population transferring from one grid cell of the current range (blue) and another grid cell within the study area (light and dark red) under future climate. (B) For standardization of the offsets obtained in step A, we first calculate spatial offsets under current climate among grid cells currently evidently occupied by red spruce (indicated by tree symbols). Next, we probabilistically re-express the raw temporal and spatio-temporal offsets as z-scores based on the empirical cumulative density function of spatial offsets (following quantile normalization, see Appendix S3: Figure S2 for more detail). For the spatio-temporal offsets only, we then apply a standardized offset threshold that categorizes all spatio-temporal offsets as either biologically tolerable (coded as 1) or non-tolerable (0) in a ‘transferability matrix’ comprising all potential donor and recipient cells. Whereas the offset standardization is not a requirement for the calculation of transferability, it helps lend a biological interpretation to the offsets and the identified thresholds. (C) Standardized Local Offsets (LO) are simply the z-scores of the temporal offsets from step A and estimate disruption of existing genotype-environment associations (see Figure 2CD for LO maps). Recipient Importance (RI) of each cell in the study area is calculated as the percentage of donor cells from which immigration is possible without exceeding the tolerable offset threshold (see Figure 4CD for RI maps). RI can be interpreted as a climate suitability metric that accounts for intra-specific variation in climate adaptation. Donor Importance (DI) of a grid cell in the red spruce range is computed as the percentage of recipient cells to which emigration is possible without exceeding the tolerable offset threshold (see Figure 5 for DI maps). Note that both RI and DI can be calculated for subsets of donor and / or recipient cells based on the same underlying model by sub-setting the transferability matrix before calculating the percentages (see Figures 5 (subsets of recipients) and 6-7 (subsets of donors) for examples).

In this study, we provide an efficient way to standardize genomic offset metrics and propose the concepts of donor and recipient importance based on these standardized offsets. We test the relevance of these developments using the forest tree red spruce (*Picea rubens*) - an economically and ecologically important conifer species in temperate forests of the northeastern United States and adjacent Canada. Red spruce is anticipated to face high risk from climate change (Hamburg and Cogbill 1988; Ribbons 2014; Taylor et al. 2017) and significant loss of suitable habitat is predicted by ENMs, especially for the United States (Beane and Rentch 2015; Capblancq et al. in revision; Peters et al. 2019; Koo, Patten, and Madden 2015). Several recent studies have demonstrated local adaptation to climate in red spruce (Butnor et al. 2019; Capblancq et al. 2023; Prakash et al. 2022), and established that genetically distinct populations occur near the fragmented southern range edge (Capblancq, Butnor, et al. 2020), which may serve as crucial reservoirs for pre-adaptations to future climates.

To estimate the potential future impact of G - E disruption in red spruce, we standardized genomic offsets relative to contemporary adaptive genomic variation across the landscape and evaluated their predictive power using growth performance data from red spruce seedlings grown in three common gardens. We then defined a standardized offset threshold assumed to be tolerable for red spruce and developed two new metrics: (1) Donor Importance (DI), defined as the percentage of recipient locations to which a given population could be transferred without exceeding a threshold level of biologically tolerable genomic offset, and (2) Recipient Importance (RI), defined as the percentage of donor populations that could be transferred to a given location, each without exceeding the tolerable offset threshold. These metrics allow the estimation of transferability across the landscape while minimizing G - E disruption. We applied our offset-based approach in combination with classical ENMs to address the following questions:

1. How do assessments of future climate suitability from species-level ENMs compare to climate change risks estimated using offset-based metrics (LO, DI, and RI) that take local adaptation into account?
2. How do prognoses for local persistence and the need for interventions through assisted migration differ between geographically and genetically distinct groups of populations?

## Material and Methods

### Study species and field sampling

Red spruce (*Picea rubens* Sarg.) thrives in cool and moist conditions with a range that extends along the spine of the Appalachians and into eastern Canadian maritime provinces, southern Québec and eastern Ontario. In the Central and Southern Appalachians, red spruce populations occur mainly as fragmented mountaintop “sky islands” (Little 1971) (see Figure 2A for current range prediction). Red spruce represents a keystone species for the conservation of vulnerable plant and animal species, as well as the capacity of mixed red spruce and northern hardwood forests to act as carbon sinks (Clipp et al. 2022; Rentch et al. 2016). Intensive logging and subsequent fires in the 19th and early 20th centuries (Adams and Stephenson 1989), followed by air pollution later in the 20th century substantially reduced the distribution of red spruce (Johnson and Siccama 1983; Mathias and Thomas 2018; Verrico et al. 2020). We therefore lack a clear picture of its natural realized niche, although scattered occurrences may still approximate the extent of occurrence before colonization (e.g., see Byers et al. (2013) for West Virginia).

**Figure 2:**
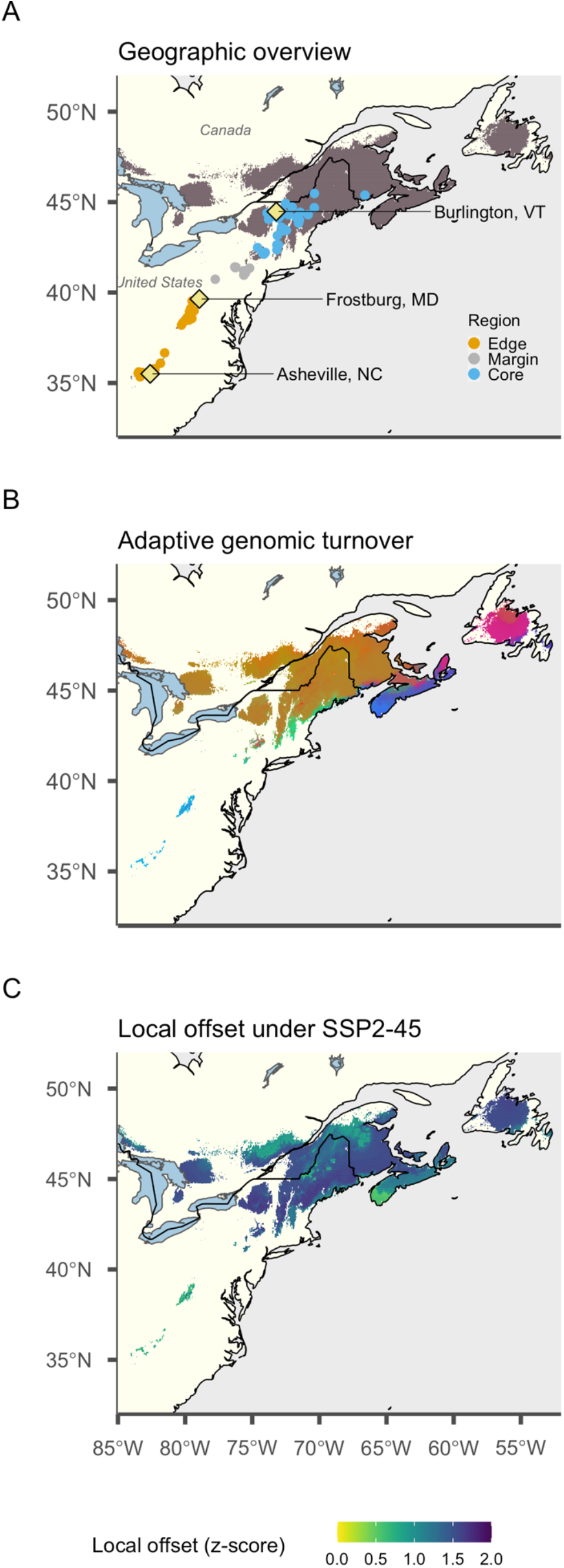
Overview of our study design in relation to red spruce’s (*Picea rubens*) current predicted distribution, climate-adaptive genomic differentiation and expected climate change disruption. (A) Current predicted distribution (climate suitability >0.25) from ENMs fitted using seven climate variables (ENM predictors, Table 1). Points indicate the locations of 64 red spruce populations that comprise three different geographic regions for use in the genomics and common-garden studies. Diamonds indicate the locations of our three common gardens. (B) Spatial pattern of genetic variation at 240 candidate SNP (Single Nucleotide Polymorphism) loci for climate adaptation as predicted by a Gradient Forest model based on eleven climatic variables (GF predictors, Table 1). Grid cells colored similarly have similar predicted genomic composition. (C) Local Offsets quantify the expected disruption of those existing genotype-climate associations under a moderate shared socio-economic pathway (SSP2-45).

**Table 1:**
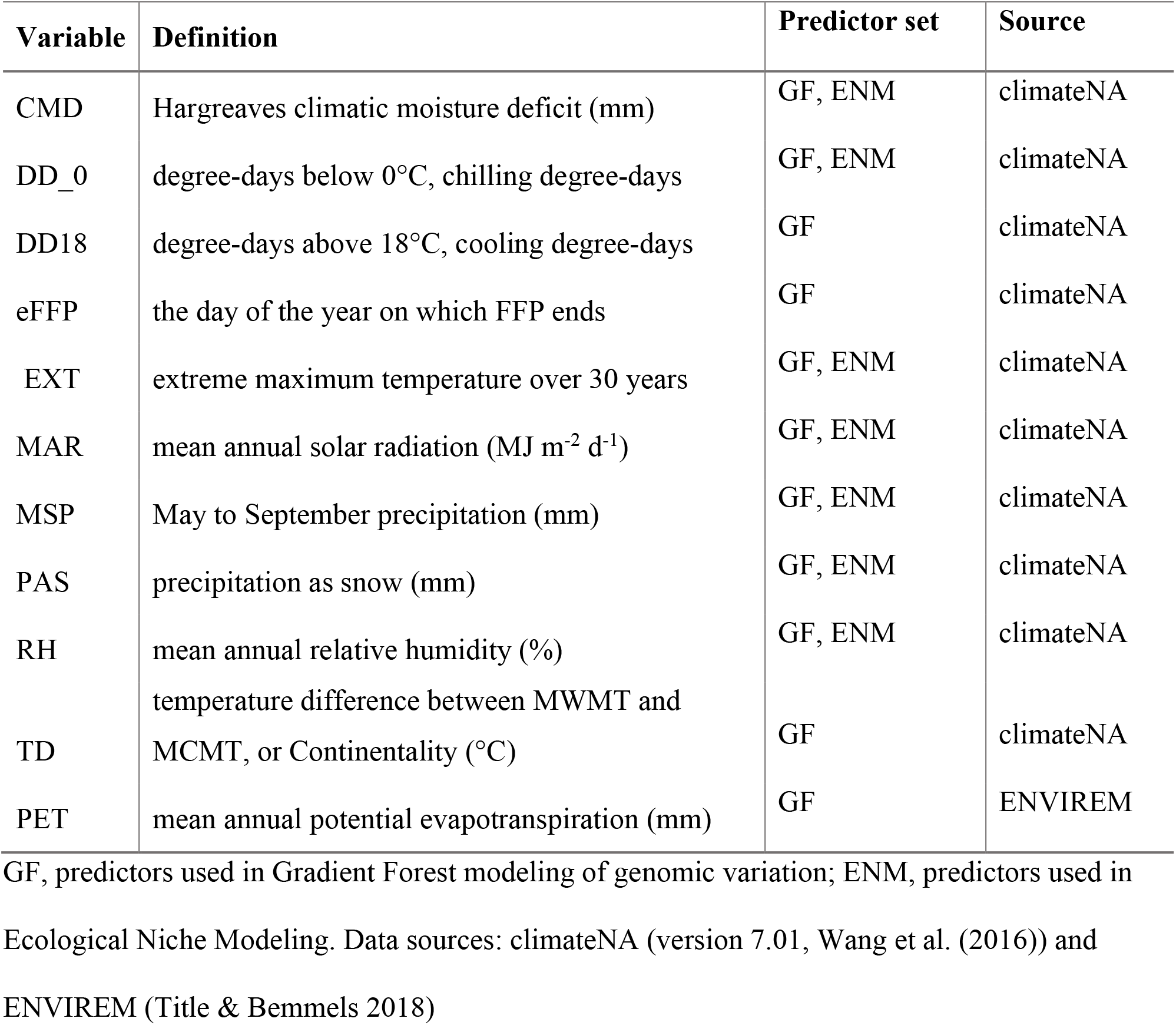
The set of eleven climatic variables used in this study

The northward range shift of red spruce from a southern refugium after the last glacial maximum (Capblancq et al. in revision) led to population genetic differentiation between the northern and southern parts of the range, and the evolution of local adaptation along geographically co-occurring climatic gradients (Capblancq et al. 2023). Using GEA approaches, Capblancq et al. (2023) identified 240 Single Nucleotide Polymorphism (SNP) candidates associated with different climatic gradients that grouped into two clusters of genes related to responses to abiotic stresses, especially drought and extreme heat. The field sampling locations for the common garden studies and genomic data utilized here (Figure 2A) span large parts of the current red spruce distribution and comprised three distinct geographic regions (the northern range *core*, the mid-latitude range *margin*, and the southern range *edge*) that have previously been shown to reflect neutral as well as adaptive genetic population structure (Capblancq, Butnor, et al. 2020; Capblancq et al. 2023).

### Climatic data for Gradient Forest and Ecological Niche Modeling

We delineated our study area to all ecoregions in which red spruce has been recorded to occur as well as adjacent ecoregions (*sensu* Dinerstein et al. (2017)) – an area that ranged from approximately -100°W to −48°W and 20°N to 60°N. To characterize climate, we used variables from the climateNA (version 7.01, Wang et al. (2016)) and ENVIREM (Title and Bemmels 2018) databases at a resolution of 2.5 arc-minutes (∼ 5 × 5 km^2^). Both the current distribution and local adaptation of long-lived tree species over generations of natural selection primarily reflect historical climatic conditions. To represent these historical conditions, we chose the standard normal period 1961-1990 CE defined by the World Meteorological Organization (WMO), which has a dense coverage of weather station data underlying the interpolated datasets and precedes the significant warming of the last ∼25 years (Wang et al. 2016).

Some of the ENM algorithms are more sensitive to collinearity of predictors than is GF, we thus used slightly different sets of variables for the two approaches. In predictor selection for GF modeling, we focused on annual variables, which were highly correlated with corresponding monthly and quarterly variables. We applied an iterative process to the annual climate data obtained for our field sampling locations with the goal of retaining a final set of the most biologically relevant and least collinear variables. To this end, we combined three different approaches: 1) assessment of collinearity using variance inflation factor (VIF), 2) characterization of the major climatic axes influencing genomic variation across the entire genomic set of candidate loci using RDA (Capblancq et al. 2023) and 3) evaluation of variable importance using preliminary GF model fitting. Details on the selected variables (hereafter, ‘GF predictors’) are summarized in Table 1. For the ENMs, we further reduced the GF predictor set of 11 to seven variables (Table 1, ‘ENM predictors”) that are ecologically relevant and do not exceed a VIF of 12 in the occurrence dataset of the ENMs. Note that we also fitted a second ENM ensemble using all GF predictors to assess the influence of predictor choice. The future climate projections for the time period 2071 – 2100 CE come from three CMIP6 scenarios for the moderate SSP2-45 shared socio-economic pathway (IPCC 2021), averaged over fifteen General Circulation models (Wang et al. 2016). Since future climate projections are not available from ENVIREM, we calculated future potential evapotranspiration using monthly temperature projections from climateNA and ENVIREM scripts (https://github.com/ptitle/envirem).

### Ecological Niche Modeling of current and future climate suitability at the species level

We used ENMs to estimate the realized environmental niche of red spruce and project changes in climate suitability before the end of the 21st century across eastern North America. We documented all technical details and underlying decisions in an ODMAP protocol (Zurell et al. 2020) (Appendix S1) and only provide a summary here. Red spruce occurrence data for the United States and Canada were gathered from forest inventories and other databases as listed in Appendix S2 and comprised a mixture of presence-only and presence-absence information. The forest inventories provided many reliable absences over most of our study area. We retained the absences for grid cells that did not contain presences and modelled the data as presence-absence. After cleaning and thinning to one occurrence record per 2.5 arc-minute grid cell, which is a standard correction method for sampling bias (Guisan, Thuiller, and Zimmermann 2017), 84,712 occurrence data points remained, including 8,825 presences and 75,887 absences. A map of the presence data points and ecoregions can be found in Appendix S3: Figure S1.

We fitted ENMs using five different algorithms, including Generalized Linear and Generalized Additive Models (GLM, GAM), Multivariate Adaptive Regression Splines (MARS), Gradient Boosting Machines (GBM) and Random Forests (RF) in R (version 4.1.2, R Core Team, 2021). Ensemble mean projections of current climate suitability as well as ensemble forecasts weighted by the AUC of each fitted model were obtained from five split-validation runs. We defined folds of training and testing data sets based on spatial blocking with systematic block selection using a range of 300 km to allow for realistic error estimation in a spatially structured environment (R package ‘blockCV’, Valavi et al. (2019)). Model evaluation was conducted based on Kappa, TSS and AUC statistics (Guisan, Thuiller, and Zimmermann 2017). Variable importance was calculated according to (Boonman et al. 2020). To delineate red spruce’s current range, we converted the predicted probability of occurrence to presence/absence using the threshold at which sensitivity equaled specificity as implemented in the R package ‘dismo’ (Hijmans et al. 2020). Equal sensitivity-specificity is a commonly used and well-performing suitability threshold that minimizes the absolute difference between sensitivity and specificity (Liu et al. 2005).

### Gradient Forest modeling and genomic offset calculation

We used Gradient Forest implemented in the ‘gradientForest’ package (Ellis, Smith, and Pitcher 2012) in R (version 4.1.2, R Core Team 2021) to model spatial and temporal variation of red spruce genomic composition. We parameterized the GF models with the eleven climatic ‘GF predictors’ for our 64 sampling locations (Figure 2A). As response variables, we used allele frequencies of 240 candidate SNP loci previously identified by Capblancq et al. (2023) as strongly associated with climatic gradients. We trained the GF models by fitting 5000 regression trees per SNP. The variable correlation threshold to invoke conditional importance estimates was set to 0.5 and default value (number predictors / 3) was used for the number of predictors randomly sampled as candidates at each split. For the calculation of model predictions we used the default setting of linear extrapolation beyond the current climatic conditions of our sampled populations.

Genomic offsets based on GF are calculated as distances between population positions in the multidimensional transformed climatic space spanned by the vectors of cumulative importance of the GF model predictors (Fitzpatrick and Keller 2015). Therefore, before calculating genomic offsets, we first created GF predictions that rescaled the current and future climatic conditions represented by the eleven predictor variables into common units of genomic variation for each grid cell in the current and future range of red spruce (hereafter referred to as “transformed climate variables”). For this study we extended the classical use of genomic offset metrics to quantifying G - E disruption following transfer to a new environment in either time or space and calculated three types of offsets (see conceptual Figure 1): a) contemporary spatial offsets were calculated among all geographic grid cells (2.5 arc minutes) currently occupied by red spruce (based on the 8,825 presences in our occurrence data set) using transformed current climate predictors; b) temporal offsets quantified the offset between current and future climate (i.e., Local Offset) in each grid cell in the current range of red spruce predicted by the ENM ensemble; c) spatio-temporal offsets describe the G - E disruption for a population being transferred from one grid cell in current climate to another grid cell in future climate.

### Standardization of genomic offsets

Genomic offsets based on GF are calculated in common units of expected genomic differentiation between places and/or times. Thus, they should better represent the expected biological impacts of climate change compared to raw climate differences uninformed by adaptive genomic information, which has been supported empirically (Capblancq and Forester 2021; Fitzpatrick et al. 2021) and using simulations (Láruson et al. 2022). However, two mathematical properties hamper the interpretability of the offset values: First, instead of a biologically-defined metric such as the average allele frequency change, GF predictions represent the variance in allele frequencies explained by environmental gradients by a given model for a given data set, so their interpretation is not translatable across studies. Second, Euclidean distances automatically increase with dimensionality, i.e., the number of transformed variables included in the distance calculation - a problem known as the *curse of dimensionality* (Bellman 1957). As such, the magnitude of genomic offsets from GF is comparable only in a relative sense (e.g., northern populations were predicted to have higher offsets than southern populations) and are not directly comparable between studies – or even between models for the same species fit with different numbers of predictors. Furthermore, GF offsets lack a direct link to the amount of maladaptation a population can tolerate. These issues have parallels with determining the magnitude of climate change that will result in a no-analogue climate, and we propose that work in this area offers a partial solution for standardizing genomic offsets to improve their biological interpretability. Mahony et al. (2017) introduced an intuitive metric termed sigma dissimilarity that quantifies the magnitude of climate change relative to past climatic variability at a location. To this end, the authors transformed Mahalanobis distances between climates in two points in time at a location into z-scores, such that they expressed as probabilities based on the distribution of local historical interannual climate variability.

Here, we adapt sigma dissimilarity for standardizing GF genomic offsets relative to contemporary genomic variation across the landscape (i.e., spatial offsets) by probabilistically re-expressing temporal offsets as z-scores of the distribution of spatial offsets. However, we decided against using Mahalanobis distances for the following reasons: (1) Mahalanobis distance assumes a multivariate normal distribution of the variables, whereas GF cumulative importance functions are always bounded between 0 and the maximum *R^2^* across all SNPs, resulting in skewed transformed climate variables; (2) The Mahalanobis distance rescales the variables before distance calculation, which in our case would remove the signature of variable importance that is critical for properly weighting the climate variables in the estimation of offsets; (3) The Mahalanobis distance removes collinearity between variables, which we wish to retain, as Gradient Forest accounts for inflation in variable importance for correlated predictors during model fitting and we assume that any remaining correlation in the transformed climate variables reflects biological significance.

We thus adjusted Mahony et al.’s (2017) approach as follows (see Appendix S3: Figure S2): (1) We first used GF to estimate spatial genomic offsets between all currently occupied grid cells; (2) We then calculated temporal and spatio-temporal GF offsets as Euclidean distances using the methods described in Gougherty et al. (2021); and (3) standardized the temporal and spatio-temporal offsets by expressing them as probabilities of an empirical distribution of all contemporary spatial offsets (i.e., the distribution of spatial genomic offsets obtained in Step 1). Finally, we (4) converted the probabilities as percentiles of a chi distribution with one degree of freedom -- an approach used by Mahony et al. (2017) and also commonly known as quantile normalization (Bolstad et al. 2003). Step 3 probabilistically scales temporal offsets relative to the present-day genomic offsets between populations across the geographic distribution of the species, while also removing the confounding effects of dimensionality (though requiring that the same GF model is used for all offset calculations, both spatial and temporal). The quantile normalization of step 4 allows probabilistic comparisons of offsets among studies based on easily interpretable z-scores.

### Evaluation of standardized genomic offsets

For assessing the extent to which our standardized genomic offsets predict declines in plant performance, we tested their effect on height growth of juvenile red spruce in three common gardens. Details on the experimental design, cultivation and monitoring are described in Prakash et al. (2022). In brief, three gardens were established along a latitudinal gradient including locations near Asheville, NC (35.504°N, −82.6°W, 655 m a.s.l.), Frostburg, MD (39.642°N, - 78.939°W, 588 m a.s.l.) and Burlington, VT (44.476°N, -73.212°W, 59 m a.s.l.). The seed families utilized here were identical to those used in the genomic analyses and GF model fitting (Figure 2A). We used populations means of family-level BLUPs (best linear unbiased predictors, see Prakash et al. (2022)) of shoot height increment between the ends of the growing seasons of 2019 and 2020. Current climate (1961-1990 CE) for the source populations and annual values averaged over 2019 and 2020 for the garden locations were derived from climateNA. Since 2019-2020 climate data were not available from ENVIREM, we used monthly temperatures from climateNA and ENVIREM scripts (https://github.com/ptitle/envirem) to calculate potential evapotranspiration. We used those same temperature data to calculate summer extreme temperature for 2019 -2020. Moreover, 2019-2020 mean annual radiation was not yet available from climateNA and therefore we calculated this variable using radiation data from GRIDMET (Abatzoglou 2013).

We fitted linear models with population mean height growth as the response and standardized genomic offset as a fixed effect for each garden separately. For comparison, we conducted the same analyses using raw Euclidean climate transfer distances that were standardized in the same way as the genomic offsets by probabilistically re-expressing them as z-scores based on the distribution of raw contemporary spatial climate distances. Here, the climate variables for all grid cells currently occupied by red spruce, our sampled red spruce populations, and the common gardens, were centered and scaled and all Euclidean distances calculated in the same PCA space.

### Calculation of transferability and Donor / Recipient Importance

After the offsets were standardized, we used a threshold to categorize them as tolerable (i.e., below threshold, coded as 1) or non-tolerable (i.e., above threshold, coded as 0). We chose an offset z-score threshold of 1 σ (sigma), which corresponds to the offset value below which ∼68% of spatial genomic offsets between current populations fall under current climate. While any such threshold is arbitrary, and will depend on the spatial scale of local adaptation and the frequency of different environments across the range, it offers an objective means for comparisons across studies. The resultant binary (tolerable/non-tolerable) matrix represents the estimated transferability of propagules between all potential donor and recipient cells (Figure 1B). Donor Importance (DI) for each donor cell was calculated as the percentage of recipient cells with a standardized offset below the z-score threshold. DI thus quantifies the representation of climates suitable for a donor cell population within the entire recipient area under consideration. Recipient Importance (RI) of each potential recipient cell represents the percentage of donor cells with a standardized offset below the z-score threshold. As such, RI can be interpreted as a climate suitability metric for a given location that accounts for intra-specific variation in climate adaptation. Both DI and RI can easily be calculated for subsets of donor and / or recipient cells based on the same underlying model, by sub-setting the transferability matrix before calculating the percentages (see Figures 5-7 for examples).

## Results

### Genomic variation and Local Offset across red spruce’s current distribution

In the GF analysis, variation in candidate SNP allele frequencies was most strongly influenced by chilling degree-days (DD_0, Appendix S3: Figures S3, S4) with an *R^2^*-weighted accuracy importance of 0.28 and an inflection point near a 700 chilling degree-days. DD_0 was followed by potential evapotranspiration (PET; importance 0.13; inflection at ∼850 mm), precipitation as snow (PAS; importance 0.12; inflection at ∼250 mm) and continentality (TD; importance 0.12; inflection at ∼25 °C). Overall, genomic variation across the landscape (Figure 2B) showed both latitudinal and longitudinal clines across red spruce’s range, with relatively steep clines towards the coast. Populations in the Central and Southern Appalachians, along the New England coast and lower-lying foothills in New Hampshire and Massachusetts, and in Nova Scotia and Newfoundland were predicted to have a distinct genomic make-up. Large parts of the range core appear relatively homogenous with some differentiation towards higher elevations. Local Offset (LO) predicted for the end of the 21^st^ century was lowest (z-score < 1 0) in the Southern and Central Appalachians and along the coasts of New England and southern Nova Scotia, and higher for the mountainous areas of New England and towards the northern range edge.

Intermediate levels (< 1.5 0) were predicted for other parts of New England, New Brunswick and Nova Scotia, with large parts of the range core reaching LO slightly above 1.5 0.

### Genomic offset validation using common garden data

Standardized genomic offsets between red spruce source populations and the common gardens ranged from 0.96 0 to 2.8 0 in the NC garden, 0.55 0 to 1.78 0 in MD and from 1.3 0 to 1.63 0 in VT, and thus exceeded our chosen tolerable offset threshold for most of our seedlings. Population mean height growth of red spruce decreased significantly with increasing standardized genomic offset in all three gardens (NC: *F*_(1,62)_ = 17.527, *p* < 0.001; MD: *F*_(1,62)_ = 47.76, *p* < 0.001; VT: *F*_(1,62)_ = 36.324, *p* < 0.001; Figure 3A). Similarly, standardized (raw) climate transfer distances negatively affected growth in each garden (NC: *F*(_1,62)_ = 22.815, *p* < 0.001; MD: *F*_(1,62)_ = 33.424, *p* < 0.001; VT: *F*_(1,62)_ = 15.208, *p* < 0.001; Figure 3B). When comparing the AIC values (Akaike’s ‘An Information Criterion’, Akaike (1974)) presented in Figure 3, we find that for MD and VT the model using standardized genomic offsets has a notably lower AIC, and thus higher explanatory power, than the regression using standardized climate transfer distance (ΔAIC, MD: 15.4; VT: 9). However, in the NC garden, the raw climate model describes growth decline somewhat better than the genomic offset model (ΔAIC, NC: 4.1). In the NC garden, mean height growth was low with 24.7 cm and decreased on average by 1.9 cm per 0 offset, for MD we estimated a mean of 30 cm and decrease of 5.6 cm and for VT a mean of 28.4 cm and decrease of 7.7 cm. In comparison, decrease in growth was less pronounced per 0 climate transfer distance in MD (-4.6 cm) and VT (-2.2 cm), but stronger than per 0 genomic offset in NC (-3.1 cm).

**Figure 3:**
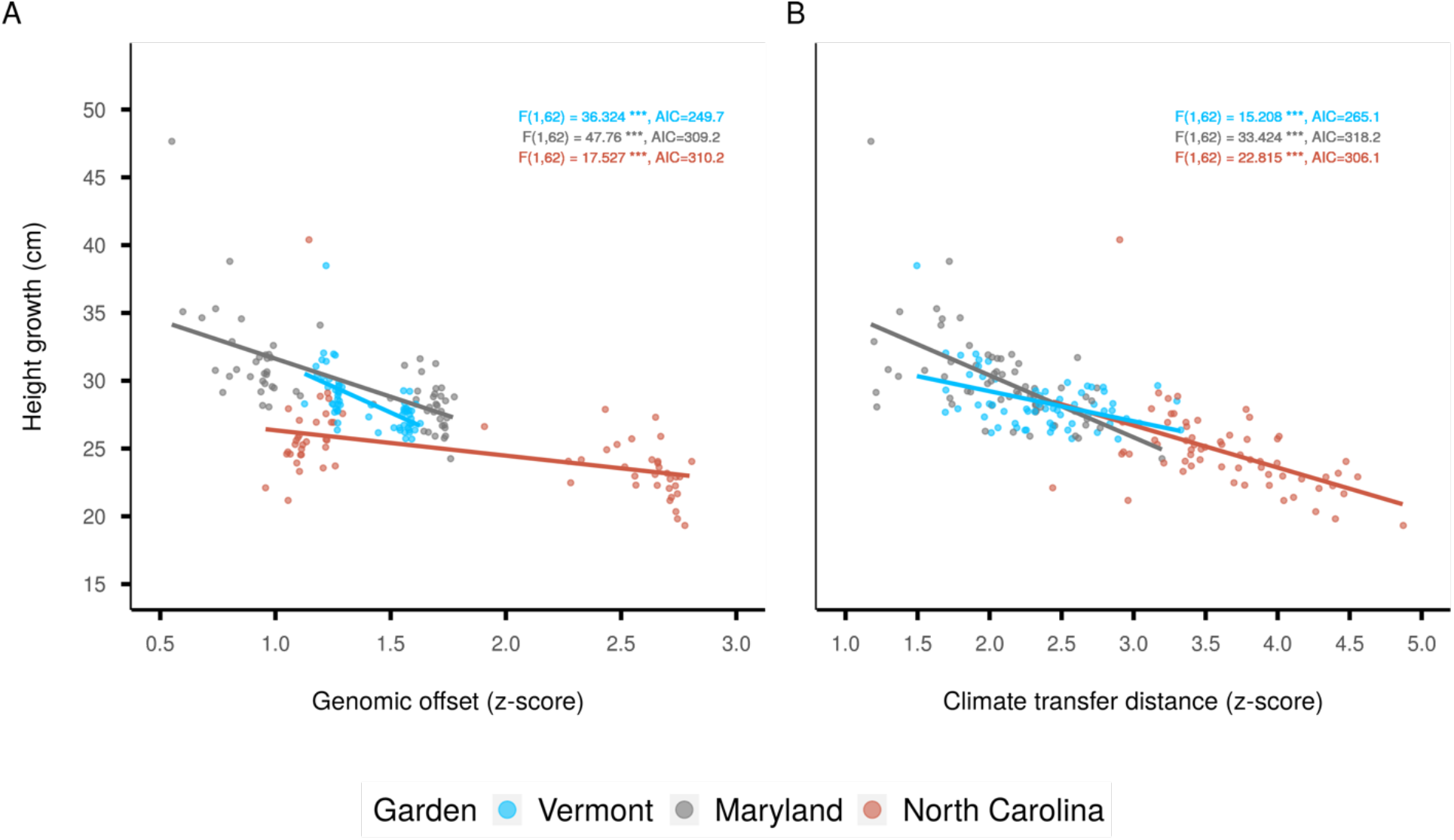
Effects of (A) standardized genomic offset predicted by Gradient Forest (GF) and (B) standardized climate transfer distance (z-score) on red spruce height growth performance (population means) in three common gardens (Figure 2A) over a 1-year period (2019-2020). Genomic offsets and climate transfer distances were calculated as Euclidean distances using current climate normals (1961-1990 CE) for the source populations and 2019-20 climate means for the gardens. Dots represent population means. F-statistics and Akaike’s ‘An Information Criterion’ (AIC) (Akaike 1974) for the separate regression analyses for each garden are given in the plots.

### ENM projections of future climate suitability and Recipient Importance

Overall, classical ENM predictions had a high accuracy (Appendix S3: Figure S6) with median Kappa per model type ranging between 0.495 and 0.520 (whole range: 0.429 – 0.749), median TSS per model type ranging between 0.740 and 0.745 (whole range: 0.665 - 0.846) and median AUC ranging between 0.921 and 0.936 (whole range: 0.910 – 0.972). Across the ENM ensemble, precipitation as snow (PAS) had the highest influence, followed by chilling degree-days (DD_0), summer extreme temperature (EXT) and climatic moisture index (CMI) (Appendix S3: Figure S7).

The weighted ensemble mean predictions of climate suitability under current climate match the distribution of occurrence data well (Appendix S4: Figure S1, Appendix S3: Figure S8) with the notable exception of an overprediction of occurrence west of Algonquin Provincial Park (ON, Canada) and in Newfoundland. But note that, whereas our occurrence data set did not contain any presences in Newfoundland, red spruce has isolated natural occurrences there (https://newfoundland-labradorflora.ca/flora/dview/?id=129). In response to future climate, the ENMs project drastic reductions of climate suitability across red spruce current range (Figures 4A), particularly in the United States.

**Figure 4:**
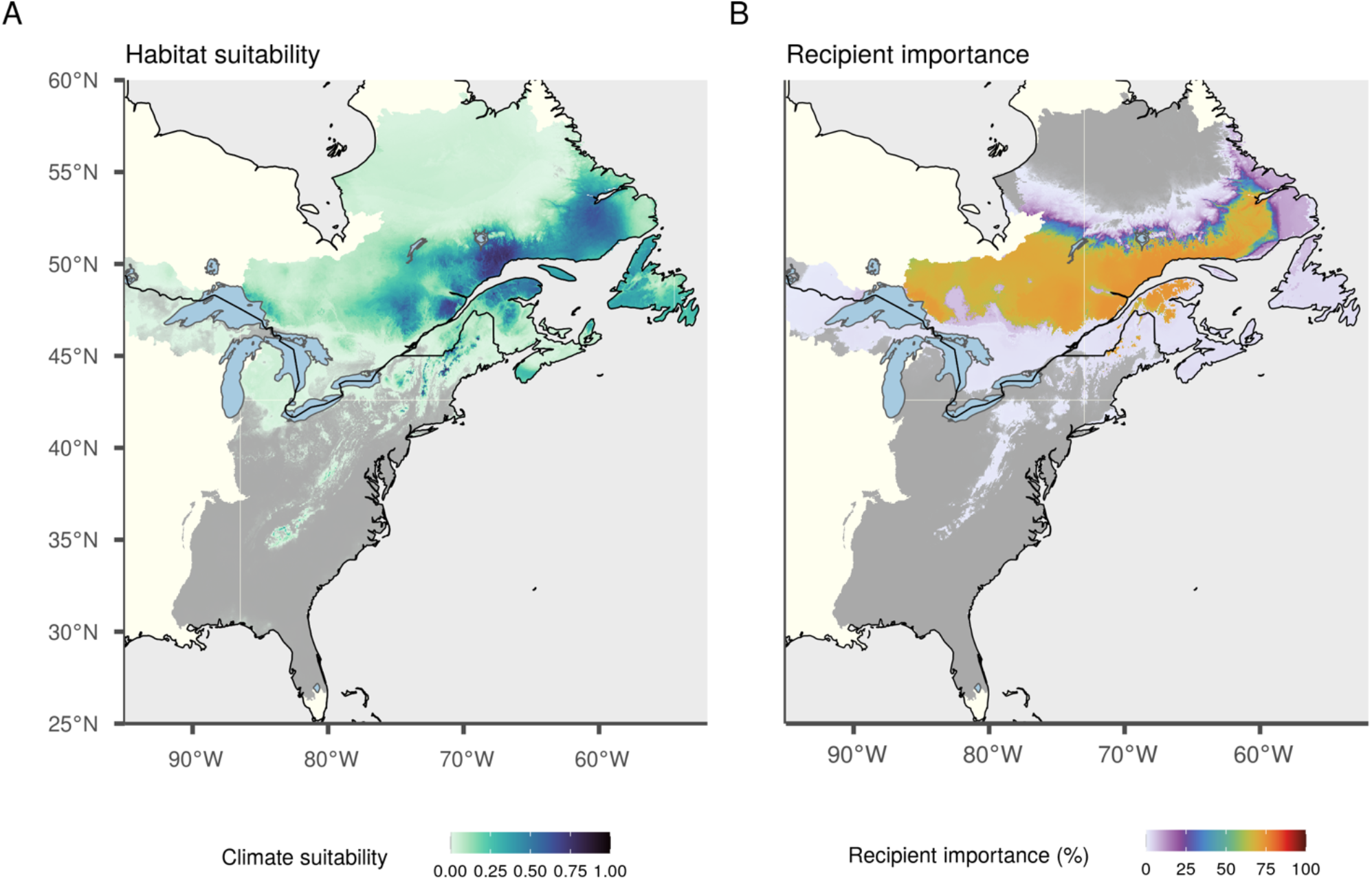
Projections of future climate suitability for *Picea rubens* by the end of the 21^st^ century assuming moderate greenhouse gas emissions (SSP2-45). (A) Weighted ensemble mean projection of red spruce occurrence probability (i.e., climate suitability) as predicted by an ensemble of ecological niche models based on seven climate variables (ENM predictor set, Table 1). (B) Recipient importance (RI) calculated based on Gradient Forest derived genomic offsets using eleven climate variables (GF predictor set, Table 1). Dark gray corresponds to near zero climate suitability / RI. ENM projections for current climate are shown in Appendix S3: Figure S8, future ENM projections using the GF predictor set can be found in Appendix S3: Figure S9.

When comparing ENM-projected climate suitability to the new metric of RI (Figure 4AB), we find broad agreement between RI and ENM-projected climate suitability, with similar trends of low RI and climate suitability in the US and high RI and climate suitability in eastern Québec and Labrador. However, areas of high RI stretch westward to regions where the ENMs project low climate suitability in the future. For results of the ENM ensemble using the GF predictor set, see Appendix S4.

### Donor Importance

Donor importance (Figure 5) expresses the suitability of a cell’s resident population to serve as a propagule donor without exceeding the tolerable offset threshold, as a percentage of the number of cells we consider as potential recipients. If considering recipients exclusively within the current red spruce range (Figure 5A), we find low DI (<15%) of the donor cells across the range with the exceptions of higher DI in the south of Nova Scotia (∼20-30%) and the southern foothills of the White Mountains (∼50%) in New Hampshire. Expanding the potential recipient area to all eco-regions currently occupied by red spruce (Figure 5B) leads to the inclusion of a higher number of recipient cells with suitable future climates as it increases DI in large parts of the range to values between 20 and 30%. Additionally including adjacent ecoregions leads to south to north and east to west gradients of increasing DI (Figure 5C).

### Genomic offset-based forecasts for geographically and genetically distinct groups of populations

To demonstrate the application of the DR and RI metrics, we explored variation among the geographically and genetically differentiated *core*, *margin* and *edge* populations as donors as well as their most relevant recipient areas (Figure 6). DI declines from the northern range *core* to the southern *edge* (Figure 6A) since we considered the entire study region as a potential recipient area (see Figure 5 for trends in DI depending on recipient area). The total suitable (i.e., non-zero RI) recipient area for our sampled populations (Figure 6A) is similar to the species-level recipient area (Figure 4B) - albeit with deviating magnitude of RI. If we assess RI separately based on sampled populations from the three geo-genetic regions (Figures 6B-D), the extent of the required northward shift increases from the range *edge* to the range *core*. Range *edge* populations are associated with high local RI in the Southern and Central Appalachians (in accordance with LO results (Figure 2C)) as well as RI values up to ∼ 50% indicating suitable future climate in isolated localities in the Adirondacks and parts of the Green and White Mountains, and large parts of Nova Scotia and Newfoundland (Figure 6B). Range *margin* populations by contrast exhibit low local RI but can find areas of high suitability (> ∼75 % RI) across the northern Appalachians, portions of the upper Midwest, and southern Canada. Range *core* populations generate above-zero RI mainly outside of the US (except some ∼100% RI fragments in Maine) in southeastern Canada along an east-west belt north of the high RI region for the range *margin* populations.

**Figure 5:**
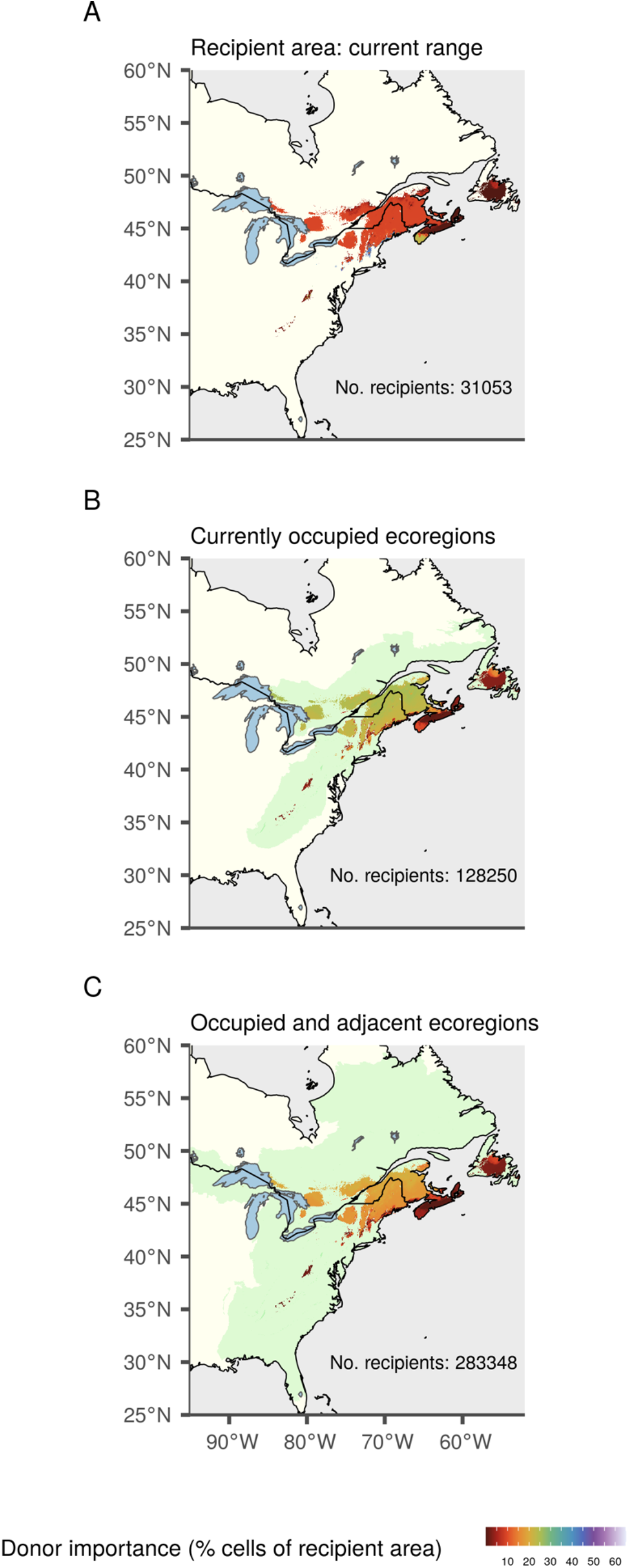
Maps of Donor Importance (DI) across the current red spruce distribution with increasing recipient area (light green) under SSP2-45. The recipient area comprises all cells in (A) the current red spruce range based on our ENM prediction, (B) ecoregions currently inhabited by red spruce, and (C) inhabited and adjacent ecoregions (ecoregions *sensu* Dinerstein et al. (2017)). Note that as recipient area increases from top to bottom, percentage DI refers to an increasing numbers of recipient cells.

**Figure 6:**
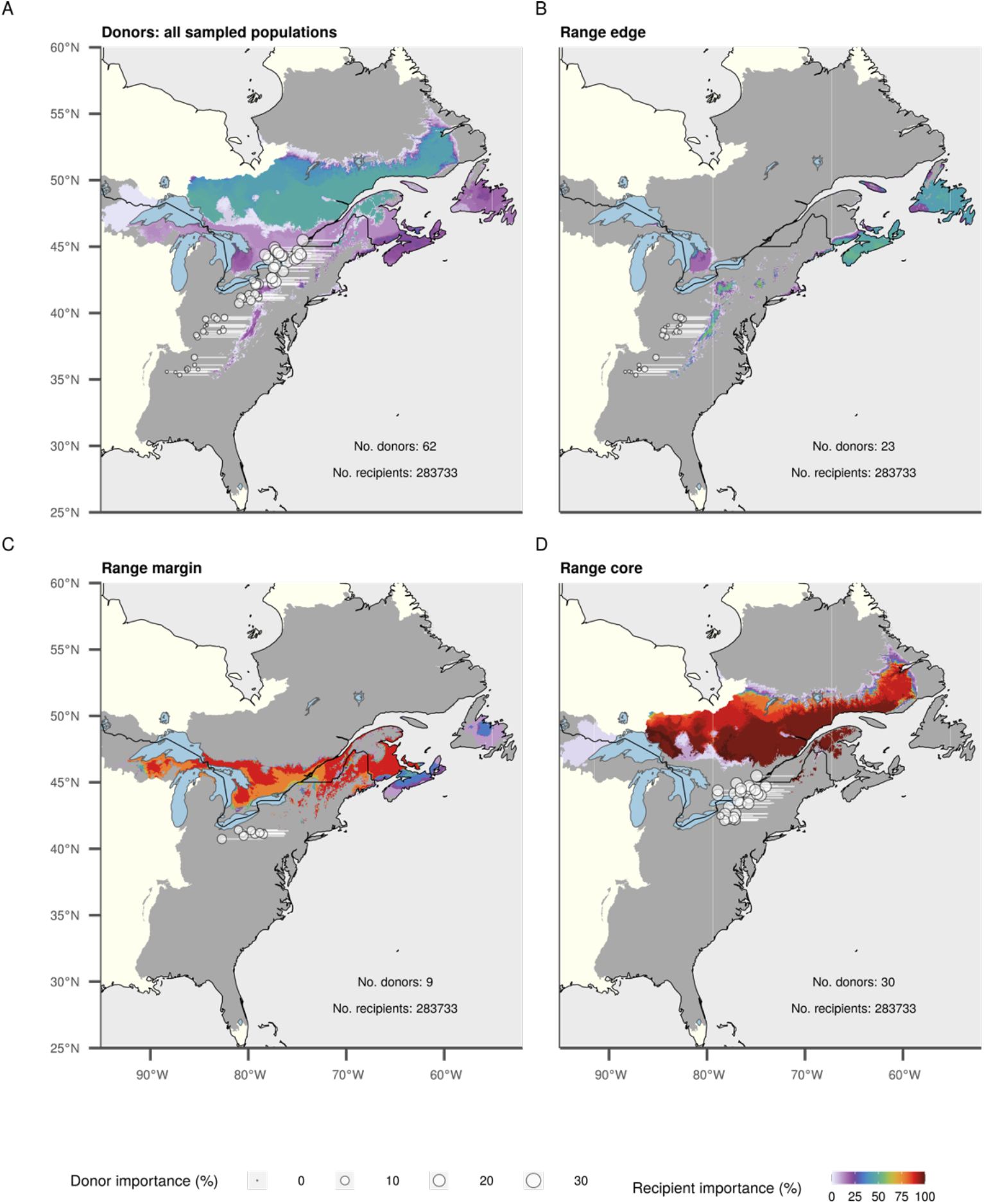
Donor and Recipient Importance considering only the red spruce populations sampled for this study (circles, see also Figure 2A) as donors under a moderate shared socio-economic pathway (SSP2-45) for 2071-2100 CE and considering the entire study region as potential recipient area. (A) All sampled populations, (B) populations of the southern range *edge*, (C) mid-latitude range *margin* and (D) northern range *core*. Please note that whereas donor importance remains constant for each population when comparing (A) with (B, C, D) and is identical to the donor importance of the respective grid cell in Figure 5C, RI of a particular grid cell changes with the number and identity of donor cells (i.e., the percentage also always refers to a different number of donors). Gray corresponds to near zero RI. A jitter and westwards shift were added to the geographic coordinates of the sampled populations; white lines point towards the actual locations.

Separating populations based on their adaptation to the most important climatic predictors in the GF model (Figure 7) reveals how adaptive differentiation along different climate variables shapes the variation in RI observed in Figure 6. For example, the drastic northward shift of suitable climate predicted for our range *core* populations seems to result from adaptations to high chilling degree-days and high precipitation as snow as well as low potential evapotranspiration and high continentality. The “RI belt” from Quebec to the Great Lakes predicted for range *margin* populations, in turn, seems to arise from adaptations to low chilling degree-days and low precipitation as snow in combination with low potential evapotranspiration and high continentality. Southern *edge* populations with adaptation to low chilling degree days, low precipitation as snow, and low continentality and low potential evapotranspiration seem to contribute most to RI in the maritime North (Figures 7F and 6B), whereas southern *edge* populations with adaptation to high potential evapotranspiration seem responsible for high local RI in southern *edge* and *margin* areas (Figures 7B and 6B).

**Figure 7:**
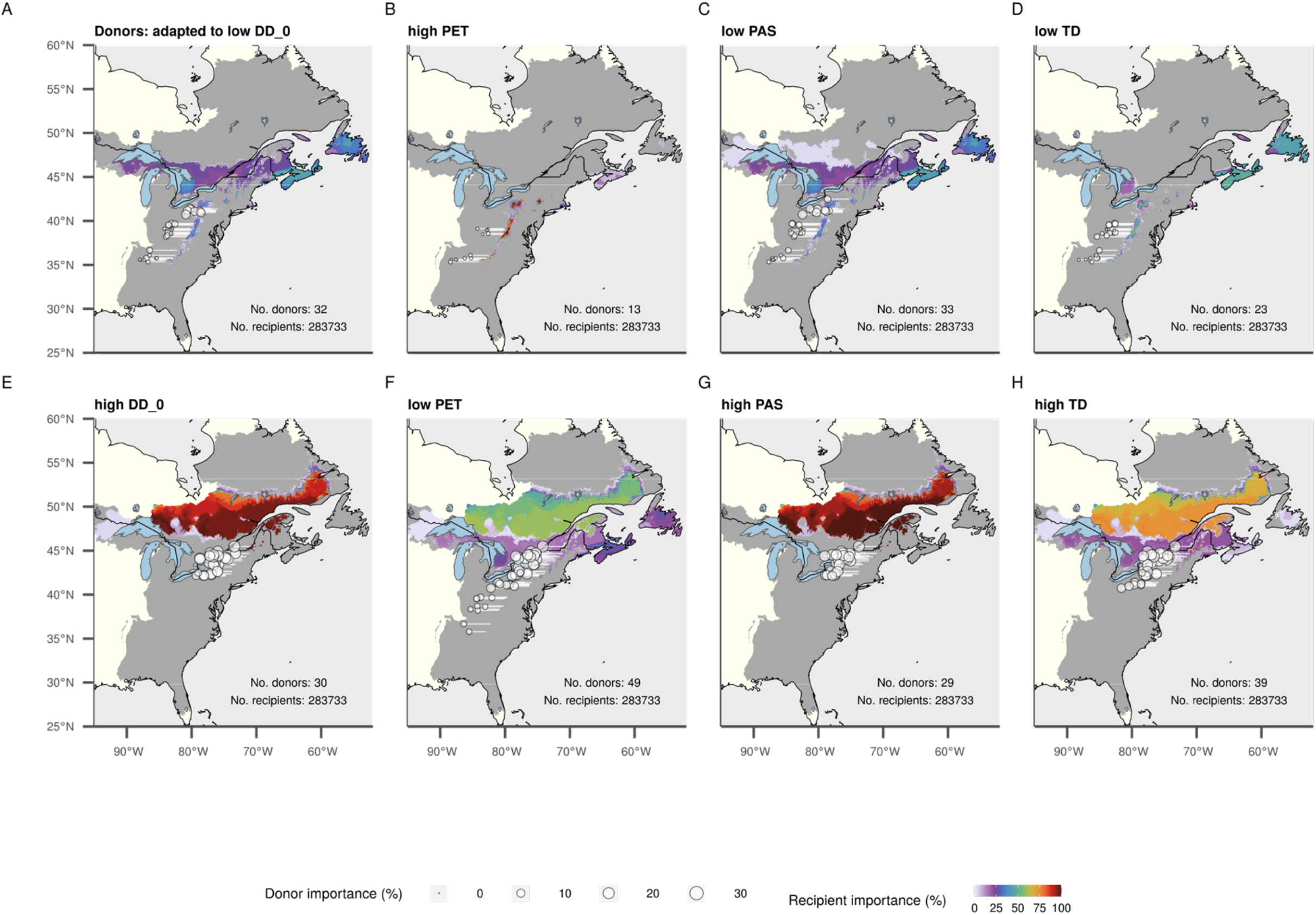
Donor and Recipient Importance (DI/RI) considering different adaptational groups of red spruce populations (circles) sampled for this study (Figure 2A) as donors under (SSP2-45 for 2071-2100 CE and considering the entire study region as potential recipient area. Donor populations locally adapted to Chilling Degree Days (DD_0) below (A) and above (E) a value of 700 mm, to Potential Evapotranspiration above (B) and below (F) a value of 850 mm, to Precipitation as Snow below (C) and above (G) a value of 250 mm, and Continentality (TD) below (D) and above (H) a value of 25 °C. Please note that DI remains constant for each population across panels and is identical to the DI of the respective grid cell in Figure 5C, RI of a particular grid cell changes with the number and identity of donor cells (i.e., the percentage also always refers to a different number of donors). Gray corresponds to near zero RI. A jitter and westwards shift were added to the geographic coordinates of the sampled populations; white lines point towards the actual locations.

## Discussion

Genomic offsets based on GF are seeing increasing application in climate change risk assessments. However, what has been lacking is an approach that allows placing the magnitude of these offsets in a more biologically meaningful context. Moreover, it is unclear how offset-based risk assessments compare with predictions from classical ENMs. Here, we developed novel standardized genomic offset-based metrics and compared them to ENM-projected climate suitability. For red spruce, ENM-and genomic offset-based forecasts were driven by similar climatic drivers and both predicted drastic shifts of suitable climate to areas north of the current range. The ENMs, however, do not provide any indication of which populations should be relocated where, whereas the offset-based metrics help assess which regions or populations may have the greatest chance for natural persistence and can guide human mitigation efforts. Here, through joint consideration of Local Offset, Recipient and Donor Importance, both for all populations collectively and for distinct groups of populations, we were able to derive refined, region-specific prognoses for potential local persistence and the need and prospects for interventions through assisted migration. Our findings suggest that few areas will be suitable for current *edge* populations under future climate and those that are suitable tend to be in proximity. In contrast, larger areas of the landscape will be suitable to most *margin* and *core* populations, but those areas are found north of current populations, suggesting a greater need for migration.

### A novel genomic offset-based approach to assessing climate change threats across species adaptive landscapes

By standardizing GF-derived genomic offsets relative to current patterns of genomic variation across the landscape, we were able to probabilistically assess the potential G - E disruption local populations might face under environmental change relative to the adaptive population differentiation currently in place across the species range. This standardization represents an advance in interpretability as compared to non-standardized GF offsets, the magnitudes of which are dependent on the number of predictors among other factors. It should be noted, however, that standardized genomic offsets still do not directly quantify changes in genetic composition (i.e., allele frequencies) and, like all ENM or GEA approaches, the inferences are constrained by the choice of environmental predictors. Our RI / DI approach extends previous applications of genomic offsets by using all potential donor - recipient matches in a study area as long as a standardized threshold is not exceeded beyond which G - E disruption is assumed to be intolerable. Here, we selected one standard deviation of contemporary spatial offsets as our tolerance threshold, which corresponds to the offset value below which ∼68% of spatial genomic offsets between current populations fall under current climate. This choice was made in view of our main objective to compare occurrence-based climate suitability predictions obtained from ENMs with offset-based metrics. Our common gardens suggested that this threshold may have been somewhat liberal, as height growth was observed to decline with increasing offset even for values up to 1.0 0, and we note that few of our gardens generated offset values <1.0 0 and none <0.5 0 (Figure 3). Depending on the application, it might be preferable to determine population-or region-specific thresholds based on criteria such as predicted reductions in performance (Wang, O’Neill, and Aitken 2010), local environmental variability (e.g., interannual climatic variability (Mahony et al. 2017), or variation in the sensitivity to disruption of local adaptation (Angert, Bontrager, and Ågren 2020; Bontrager et al. 2021; Willi and Van Buskirk 2022).

In two of our common gardens, standardized genomic offsets better predicted growth performance than did standardized climate transfer distances (Figure 3). This is in accordance with results on performance of populations of lodgepole pine (*Pinus contorta*) and balsam poplar (*Populus balsamifera*) in common gardens (Fitzpatrick et al. 2021; Capblancq and Forester 2021). Our 1 0 threshold was associated with an average decrease in growth between 1.9 and 7.7 cm depending on the garden environment. Notably, most of our planted individuals experienced a genomic offset larger than this threshold, as the gardens experienced generally warmer, drier and less seasonal climate than the multi-decadal average climates of our source populations (Prakash et al. 2022). The climate of the NC garden fell outside our sampling range along three of the most important climatic gradients (DD_0, PET, PAS; Appendix S3: Figure S4). The resulting extrapolation of the GF predictions beyond the environments used to train the models might thus have underestimated offsets for this garden, especially for the subset of populations (primarily from the *edge*) that occupy positions on the flat part of the cumulative important curve nearest to the garden location, leading to decreased explanatory performance in comparison to raw climate transfer distances. It should also be kept in mind that we have considered a fraction of the life cycle of this long-lived tree species, and thus may have missed critical stages that occur both earlier (e.g., germination) and later (e.g., reproduction) in the life history of red spruce.

### Climatic drivers of ENM-projected climate suitability and intraspecific adaptive differentiation

Climatic factors limiting a species’ niche may not necessarily be those driving intra-specific differentiation (Mahony et al. 2020). In our study, however, there was relatively high agreement in the importance of climatic variables between the ENM and GF approaches (Appendix S3: Figures S3, S7). Both analyses suggested a higher relevance of temperature than precipitation related variables, which is in accordance with results of Li et al. (2020), who analyzed provenance trials with red spruce source populations north of our sampling region. Our results also suggest that winter temperature-derived metrics such as chilling degree-days and precipitation as snow are most important for both genetic differentiation and range delineation. Possible causes underlying this finding include the low frost tolerance of red spruce compared to boreal spruce species such as *P. mariana* and *P. glauca* (Dumais and Prévost 2007), and the great importance of adaptation regarding chilling requirements and frost hardening for the optimal timing of bud-break (Dumais and Prévost 2007) and bud set (Prakash et al. 2022). Kosiba et al. (2018) further documented positive growth effects of higher fall through spring temperatures in natural red spruce stands in the Northeastern US. Ribbons (2014) found interactions of both winter and summer extreme temperatures with precipitation effects on the growth of red spruce in Massachusetts and suggested that higher temperatures increased evapotranspiration and thus negative effects of drought at some sites. This finding corresponds well with the high importance of potential evapotranspiration and extreme temperature in our study and previously reported differentiation in genes involved in acute stress response along climatic gradients characterized by extreme temperature variation (Capblancq et al. 2023). Increases in hottest yearly temperatures have also just recently been shown to be a driver of local extinctions across a wide range of plant and animal species (Román-Palacios and Wiens 2020). The low significance of May to September precipitation in our analyses was somewhat surprising, as precipitation during the growing season is considered to drive the high growth performance of red spruce in the Southern Appalachians and along the coast as compared to the rest of the range (Blum 1990). Apparently, differentiation between continental and more humid regions was better represented by the temperature based continentality metric (TD) in our GF analyses.

### ENM-and genomic offset-based forecasts support range-wide decline and poleward shifts in climate suitability

Our ENM ensemble projects climate suitability declines throughout the current red spruce range and in some regions total habitat loss (Figure 4A), which is consistent with predictions of a coupled growth and climate model for the Southern Appalachians (Koo, Patten, and Madden 2015; Koo et al. 2014) and previous ENMs for West Virginia (Beane & Rentch, 2015) and the entire United States (Peters et al. 2019). The predicted pattern of contraction is also consistent with results from a demo-genetic model by our group that fit an ENM ensemble to a different and coarser set of climate variables and incorporated population size and dispersal using population genetic variation (Capblancq et al. in revision). Among all GF-derived metrics, RI is most comparable to ENM-projected climate suitability and indeed shows similar trends throughout the current red spruce range (Figure 4B). Both RI and ENMs also predict a drastic northward shift of suitable climate. However, RI suggests higher suitability in Ontario than the ENM projections.

### Local Offset predictions suggest possible persistence at range edges

LO allowed us to quantify the expected future G - E disruption that existing populations may be exposed to *in situ*. For red spruce, comparably low LO were predicted for the southern, eastern and northern range edges (Figure 2C). The low LO at the northern range edge coincides with ENM-projected high climate suitability and high RI, especially for range *core* donor populations (Figure 6D). However, predicted GF offsets at the northern range edge need to be interpreted with caution, since we did not sample in this area and the extrapolation applied by GF may not accurately represent genomic differentiation in those areas. Further, our GF analyses assume current adaptation between population allele frequencies and the environment (Capblancq, Fitzpatrick, et al. 2020), which might not always be the case if some populations are mal-adapted under current climates and might therefore actually benefit from climatic changes. Some evidence of current climate mal-adaptation of red spruce’s northern range edge exists, for example from provenance trial experiments showing that some northern source populations benefit from transfer into warmer test site climates (Li et al. 2020). Such mal-adaptation at the leading range edge is common and might be explained by limits on adaptation due to expansion history such as population age, bottlenecks and genetic drift, or expansion load, as well as current constraints on adaptation at range edges such as genetic swamping from the core of the range (Angert, Bontrager, and Ågren 2020; Bontrager et al. 2021).

Southern *edge* populations in the Central and Southern Appalachians exhibited the lowest LO within the current spruce range, even staying below 1 0, suggesting that these populations might be able to persist *in situ*, which seems contrary not only to projections of decreasing – or total loss of - climate suitability from ENMs, but also the low species level DI and RI predicted for this region. The explanation of this apparent contradiction may lie in strong local adaptation often occurring in both climatically marginal populations that experience strong selection pressure as well as at the equatorial edges of species ranges, where local adaptation has had longer time to develop and where populations are more isolated (Bontrager et al., 2021). While in our case, range *margin* and *core* populations have higher genetic diversity than the southern *edge* due to larger effective population sizes (*N*e) and likely hybridization with black spruce (Capblancq, Butnor, et al. 2020), other conditions for strong local adaptation are met in the Central and Southern Appalachians and the mapped spatial pattern of genetic variation” (Figure 2B) as well as results on adaptive differentiation among our populations presented by Capblancq et al. (2023) support the occurrence of adaptive genomic differentiation towards the southern *edge*. However, we cannot entirely rule out the possibility that our low offset estimates in the range *edge*, which contrast with the numerous published ENM-based projections of loss of climate suitability in this region, may be an artefact associated with the extrapolation of models to future climate conditions outside the climatic space covered by our sampled populations (Appendix S3: Figure S5).

Nevertheless, the ENM-projected low climate suitability may simply reflect the marginality of future southern climates compared to the climates currently inhabited by most red spruce populations, which were heavily decimated at lower elevations by logging and fire in the 19-20th centuries (Adams and Stephenson 1989). Moreover, we do not expect a southward trend in climate suitability from the *core* towards the southern *edge* under climate warming, which explains the low predicted RI in the South. The low southern *edge* DI indicates that while specific adaptations may foster *in situ* persistence, the adaptation of southern *edge* populations to the unique climate of the region might not be suitable to colonize large areas further north even under climate warming, although some areas of the current range might be candidates for assisted gene flow from southern *edge* populations.

### Implications for in situ conservation vs. assisted migration

Despite the cautiously optimistic perspective of potential persistence at the southern *edge*, the synthesis of range-wide ENM-projected climate suitability, RI and LO suggests that most existing red spruce populations will experience notable decline in performance at their current locations. Thus, assisted migration, as previously suggested for red spruce, e.g., by Byers and Norris (2011), may become necessary to protect substantial parts of the gene pool. Range-wide maps of DI (Figure 5) that consider recipient areas of varying extent highlight the need for assisted migration beyond the current range, as transferability is very low within the current range under the projected climate change (most populations could transfer to ∼10% of the current range). For current range *margin* and *core* populations, we predict high LO and near zero RI within the region but very high RI at the northern range edge and beyond, which suggests that these populations might strongly depend on assisted northwards migration beyond the current range. For southern *edge* populations, the region-specific DI and RI analyses (Figure 6) raise the possibility of *in situ* persistence in some cases for local populations (small LO) but also more broadly within the region under local dispersal or assisted migration. The results also imply that Central and Southern Appalachian populations could serve as donors for assisted migration into the northern Appalachians as well as the Canadian Maritime provinces.

Separating our sampled populations into climate-based adaptational groups (Figure 7) allowed us to further interpret the geographic region-specific results (Figure 6) by examining how specific climate adaptations shape the populations’ potential to serve as donors for various recipient locations in our study area. Thus, these analyses also reveal which adaptations render populations either most suitable for *in situ* conservation, such as high potential evapotranspiration and high continentality populations in the Southern and Central Appalachians or which populations might be dependent on assisted migration, such as range *core* populations adapted to high chilling degree days and high precipitation as snow or range *margin* and *edge* populations with adaptations to low potential evapotranspiration and high continentality. Most likely owing to local topography, both low and high potential evapotranspiration populations occur in close vicinity in southern *edge* and *margin* regions, suggesting that seed sourcing schemes within this region should take this into consideration. Thus, to support conservation planning beyond our sampled populations, the SNP loci with the highest accuracy importance and linked to relevant functional genes for these climate predictors Capblancq et al. (2023) could be used as genetic markers to determine the value of targeted populations for assisted migration programs or to evaluate the expected success of *in situ* conservation for red spruce populations on the ground. Again, some caution and further research is advisable where projected future climate exceeds our sampled climatic ranges, which is the case for four of our high PET range *edge* populations under future climate (Appendix S3: Figure S5). It is also important to consider other risk factors in conservation planning, such as inbreeding depression, which has been demonstrated in southern *edge* red spruce populations (Capblancq et al. 2021). Therefore, even for *in situ* conservation it might be advisable to induce admixture among local populations that exhibit adequate climate adaptations in efforts to maximize the available locally adapted genetic diversity (Potter et al. 2017).

## Conclusions

Landscape and evolutionary genomics continue to gain attention in global change biology and conservation science (Isabel, Holliday, and Aitken 2020; Mahony et al. 2020; Sork et al. 2013; Waldvogel et al. 2020) and particularly in the use of genomic offset metrics (Capblancq, Fitzpatrick, et al. 2020; Dauphin et al. 2021; Gougherty, Keller, and Fitzpatrick 2021; Nielsen et al. 2021; Rellstab, Dauphin, and Exposito-Alonso 2021). Our approach extends previous applications of the genomic offset concept as it constrains donor - recipient matches relative to the contemporary genomic turnover across a species range. This standardization should facilitate management applications such as seed source and planting site selection for assisted migration that takes advantage of the genetic variation in climate adaptations present among the populations of a species. Moreover, the GF population-level approach may be preferable over ENMs as GF does not require the ability to estimate the full realized niche, which is useful for species such as red spruce for which the natural distribution is unknown. The full strength of the DI-RI approach to recognizing and considering local adaptations in forecasting and management planning becomes apparent when we consider groups of potential donors separately *based on the same underlying model*. In our case, this approach identified the southern *edge* populations as best candidates for *in situ* restoration and assisted gene flow (within the current range), but strongly advocates for the consideration of assisted colonization northward (beyond the current range) for range *margin* and *core* populations. Subsets of the transferability matrix can be selected based on any kind of ecological, geographic or genetic criteria for donors and recipients alike, allowing for applications beyond those in this study. A critical aspect for further developments of standardized genomic offsets will be the selection of tolerable thresholds and the use of sensitivity analyses to explore how these and other decisions impact forecasts and associated management implications.

## Acknowledgements

This project was supported by awards from the National Science Foundation (1656099 and 1655344), a USDA-HATCH award (1006810), and a UVM Postdoctoral Fellowship award. We appreciate the assistance of many individuals who contributed to the collecting of genetic samples and assisted with the establishment of and data collection at the common gardens, including Jacquelyne Adams, John Butnor, Sonia DeYoung, Erica Duda, Natalie Haydt, Kurt Johnsen, Matthew Lisk, Helena Munson, David Nelson, Robin Paulman, Anoob Prakash, Brittany Verrico, and Ethan Thibault. Coline Boonman shared code for the calculation of variable importance.

## Author Contributions

Matthew C. Fitzpatrick and Stephen R. Keller developed the project and obtained funding, with additional funding acquired by Thibaut Capblancq and Stephen R. Keller. All authors participated in the collection of common garden data, which were managed by Anoob Prakash and Stephen R Keller. Thibaut Capblancq conducted the genomic data analyses with advice from Stephen R. Keller. Susanne Lachmuth and Matthew C. Fitzpatrick developed the concept of standardized offsets and DI/RI metrics. Susanne Lachmuth acquired the species occurrence and climate data, performed the modeling and data analysis and drafted the manuscript with advice from Matthew C. Fitzpatrick. All authors contributed to and approved the final manuscript.

