## Appendix S1 for "Novel genomic offset metrics account for local adaptation in climate suitability forecasts and inform assisted migration"

### Novel genomic offset metrics account for local adaptation in climate suitability forecasts and inform assisted migration

#### – ODMAP Protocol –

Susanne Lachmuth, Thibaut Capblancq, Anoob Prakash, Stephen R. Keller, Matthew C. Fitzpatrick

2022-08-11

---

#### Overview

##### Authorship

Contact :

Study link: Publication in prep.

##### Model objective

Model objective: Forecast and transfer

Target output: Continuous climate suitability index, binary suitable vs. unsuitable habitat projection

##### Focal Taxon

Focal Taxon: Picea rubens Sarg.

##### Location

Location: Eastern North America (United States and Canada)

##### Scale of Analysis

Spatial extent: -100, -48, 20, 60 (xmin, xmax, ymin, ymax)

Spatial resolution: 4.625 x 4.625 (2.5 arc minutes)

Temporal extent: Occurrence data extend from 1850 to present, climate data extend from 1961 to 2100 CE.

Temporal resolution: We used 1961-1990 normals for contemporary climate and forecasts of future climate normal for 2071-2100.

Boundary: natural

##### Biodiversity data

Observation type: field survey, standardised monitoring data, citizen science

Response data type: presence/absence

#### Predictors

Predictor types: climatic

#### Hypotheses

Hypotheses: The geographic distribution of tree species in temperate climate zones is determined by growing season temperatures, climatic water balance, and seasonality (Metzger et al. 2013). Accordingly, we chose ecologically meaningful predictor variables that contemporarily show considerable variation and low collinearity across the study region and have a high potential for delineating the range limits of Picea rubens.

Reference: Metzger, M.J., Bunce, R.G.H., Jongman, R.H.G., Sayre, R., Trabucco, A. & Zomer, R. (2013). A high-resolution bioclimate map of the world: a unifying framework for global biodiversity research and monitoring. Global Ecology and Biogeography, 22, 630–638.

#### Assumptions

Model assumptions: - Red spruce distribution is in pseudo-equilibrium with the environment - Occurrence data are free from observational bias - Each occurrence record represents an independent observation - Key predictors are incorporated in the model - Predictors are estimated without error - No spatial auto-correlation of residuals - Red spruce retains its niche across space and time - Relationships fitted under current conditions apply when transferring predictions, even when projected beyond the range of the training data; no change in correlation structure of environmental variables; no change in key limiting processes

#### Algorithms

Modelling techniques: glm, gam, mars, brt, randomForest

Model complexity: The model settings were chosen to yield relatively simple response surfaces because we attempted forecasting to the end of the 21st century. The species may not be at equilibrium with the environment and non-analogous climates have been projected for some geographic regions. We allowed quadratic relationships in GLMs and GAMs, and restricted MARS, BRT and random forest models in a way to avoid overfitting.

Model averaging: We combined the five types of algorithms in a weighted ensemble approach.

#### Workflow

Model workflow: We intended to chose ecologically meaningful climate variables and used variance inflation (VIF) to avoid high collinearity among predictors. With seven selected climatic variables, we fit ecological niche models using five different algorithms (GLM, GAM, MARS, RF, GBM,). Ensemble mean projections of current climate suitability as well as ensemble forecasts weighted by the AUC of each fitted model were obtained from five split-validation runs. We defined folds of training and testing data sets based on spatial blocking with systematic block selection using a range of 300 km to allow for realistic error estimation in a spatially structured environment. Model evaluation was conducted based on Kappa, TSS and AUC statistics. We estimated variable importance and produced partial response curves and map displays for visualization and assessment of ecological plausibility.

#### Software

Software: Software: R (version 4.1.2, Platform: x86_64-pc-linux-gnu (64-bit), R Core Team (2021). R: A language and environment for statistical computing. R Foundation for Statistical Computing, Vienna, Austria. URL https://www.R-project.org/)

Code availability: Code is available here: TBA

Data availability: Data are available here: TBA

#### Data

##### Biodiversity data

Taxon names: Picea rubens Sarg. (Pinaceae)

Taxonomic reference system: Catalogue of Life: 2020-04-16 Beta

Ecological level: species

Data sources: We collected Picea rubens occurrence data from various sources including from the Botanical Information and Ecology Network – BIEN , the Biodiversity Information Serving Our Nation (BISON) database, the Natural Heritage Vegetation Database for West Virginia, the Global Biodiversity Information Facility, USDA (https://data.nal.usda.gov/dataset/witness-trees-monongahela-national-forest-1752-1899), as well as US and Canadian forest inventory programs to complement our own sampling locations. Details and references for the downloaded data sets are given in Supplement 2.

Sampling design: Random

Sample size: Prevalence / sample size after thinning: 84712 / 75887

Clipping: Ecoregions currently harboring red spruce populations as well as adjacent ecoregions (see https://ecoregions.appspot.com, Dinerstein et al. 2017) : Allegheny Highlands forests, Appalachian Blue Ridge forests, Appalachian mixed mesophytic forests, Appalachian Piedmont forests, Atlantic coastal pine barrens, Central Canadian Shield forests, Eastern Canadian Forest-Boreal transition, Eastern Canadian forests, Eastern Canadian Shield taiga, Eastern Great Lakes lowland forests, Gulf and St. Lawrence lowland forests, Interior Plateau US Hardwood forests, Mid-Atlantic US coastal savannas, New England-Acadian forests, Northeast US Coastal forests, Southeast US conifer savannas, Southern Great Lakes forests, Western Great Lakes forests

Reference: Dinerstein, E.; Olson, D.; Joshi, A.; Vynne, C.; Burgess, N. D.; Wikramanayake, E.; Hahn, N.; Palminteri, S.; Hedao, P.; Noss, R.; Hansen, M.; Locke, H.; Ellis, E. C.; Jones, B.; Barber, C. V.; Hayes, R.; Kormos, C.; Martin, V.; Crist, E.; Sechrest, W.; Price, L.; Baillie, J. E. M.; Weeden, D.; Suckling, K.; Davis, C.; Sizer, N.; Moore, R.; Thau, D.; Birch, T.; Potapov, P.; Turubanova, S.; Tyukavina, A.; de Souza, N.; Pintea, L.; Brito, J. C.; Llewellyn, O. A.; Miller, A. G.; Patzelt, A.; Ghazanfar, S. A.; Timberlake, J.; Klöser, H.; Shennan-Farpón, Y.; Kindt, R.; Lillesø, J.-P. B.; van Breugel, P.; Graudal, L.; Voge, M.; Al-Shammari, K. F.; Saleem, M. An Ecoregion-Based Approach to Protecting Half the Terrestrial Realm. BioScience 2017, 67 (6), 534–545. https://doi.org/10.1093/biosci/bix014.

Scaling: Raw data were thinned to one record per 2.5 arcminute grid cell.

Cleaning: We excluded outlier data points from federal states in which the species is not reported to occur based on the "Flora of North America / Little", photo plots from the Canadian forest inventory program as well as data types of the following categories: "unknown", "fossil", "living specimen" (e.g. in botanical gardens). Data were than thinned to the resolution of the climate data (2.5 arcminutes), which lead to the removal duplicates. Grid cells were set to 'present' as soon as a presence had been reported anywhere within the cell. Grid cells occupied by water bodies were removed.

Absence data: Non-observation of species in US and Canadian forest inventory programs was treated as absence.

Background data: Not applicable.

#### Data partitioning

Training data: We defined five folds of training and testing data sets based on spatial blocking with systematic block selection using a range of 300 km to allow for realistic error estimation in a spatially structured environment

Validation data: We defined five folds of training and testing data sets based on spatial blocking with systematic block selection using a range of 300 km to allow for realistic error estimation in a spatially structured environment

Test data: N/A

#### Predictor variables

Predictor variables: We used two sets of seven and eleven climate variables downloaded from climateNA (version 6.40, Wang et al. 2016) and ENVIREM (Title et al. 2018). Details on this variable set can be found in Table 1 (main text).

Data sources: Data sources: climateNA (version 6.40, available from https://sites.ualberta.ca/~ahamann/data/climatena.html, accessed 01/28/2021, Wang et al. 2016) and ENVIREM (Avaiable from: https://envirem.github.io/, accessed 01/29/2021, Title et al. 2018)

References: Title, P.O. & Bemmels, J.B. (2018). ENVIREM: an expanded set of bioclimatic and topographic variables increases flexibility and improves performance of ecological niche modeling. Ecography, 41, 291–307. Wang, T., Hamann, A., Spittlehouse, D. & Carroll, C. (2016). Locally Downscaled and Spatially Customizable Climate Data for Historical and Future Periods for North America. PLOS ONE, 11, e0156720.

Spatial extent: -100, -48, 20, 60 (xmin, xmax, ymin, ymax)

Spatial resolution: 2.5 arcminutes

Coordinate reference system: +proj=longlat +datum=WGS84 +no_defs +ellps=WGS84 +towgs84=0,0,0

Temporal extent: 1961-1990

Temporal resolution: Means of standard normal period defined by the World Meteorological Organization (WMO) 1961-1990

Data processing: See original publications: Title, P.O. & Bemmels, J.B. (2018). ENVIREM: an expanded set of bioclimatic and topographic variables increases flexibility and improves performance of ecological niche modeling. Ecography, 41, 291–307. Wang, T., Hamann, A., Spittlehouse, D. & Carroll, C. (2016). Locally Downscaled and Spatially Customizable Climate Data for Historical and Future Periods for North America. PLOS ONE, 11, e0156720.

Errors and biases: See original publications: Title, P.O. & Bemmels, J.B. (2018). ENVIREM: an expanded set of bioclimatic and topographic variables increases flexibility and improves performance of ecological niche modeling. Ecography, 41, 291–307. Wang, T., Hamann, A., Spittlehouse, D. & Carroll, C. (2016). Locally Downscaled and Spatially Customizable Climate Data for Historical and Future Periods for North America. PLOS ONE, 11, e0156720.

Dimension reduction: For variable selection, we applied an iterative process to climate data obtained for our genomic sampling locations (see main text) with the goal of retaining a final set of the most biologically relevant and least collinear variables. To this end, we combined three different approaches: 1) assessment of collinearity using variance inflation factor (VIF) analysis, 2) characterization of the major climatic axes influencing genomic variation across the entire genomic marker a set using RDA and 3) evaluation of variable importance using Gradient Forest model fitting. Details on the selected variables are summarized in Table 1 (main text).

#### Transfer data

<Data sources>

Spatial extent: -100, -48, 20, 60 (xmin, xmax, ymin, ymax)

Spatial resolution: 2.5 arc minutes

<Temporal extent>

<Models and scenarios>

#### Model

##### Variable pre-selection

Variable pre-selection: Among the available climateNA variables we focused on the annual variables, which we complemented with annual PET from ENVIREM.

##### Multicollinearity

Multicollinearity: The selection of variables was tailored to the identification of selected loci and genomic offset calculation using Gradient Forest. The latter is not sensitive to the inclusion of correlated variables. We thus used data for our genomic sampling locations and applied a rather moderate elimination of correlated variables. We first applied a step-wise variable elimination based on the Variance Inflation Factor (vifstep, R package usdm) with a moderate VIF threshold of 20, which yielded nine variables with a VIF < 18: CMD (VIF 3.18), DD_0 (17.69), DD18 (9.71), EXT (9.34), MAR (3.74), MSP (2.99), PAS (11.9), PET (12.32), RH (3.03). We complemented this variable set with two further variables (TD, eFFP) that were of high importance in preliminary GF analyses and aligned with the two major climatic axes influencing genomic variation across the entire genomic marker a set in a preliminary RDA.

##### Model settings

glm: family (binomial), formula (makeFormul (makeFormula("PresAbs",calib[,predictors],"quadratic",interaction.level=0) ), notes (stepwise model selection based on AIC (both directions), linear terms were not allowed to drop out)

gam: family (binomial), smoothTerms (s_smoother (2, 3) ), method (GCV.Cp), select (FALSE), notes (stepwise model selection based on AIC (both directions), linear terms were not allowed to drop out)

mars: formula (~ 1+ DD_0 + DD18 + MSP + NFFD + eFFP + PAS + RH + TD + EXT + CMD + MAR +PET), degree (1), penalty (2), nk (default: min(200, max(20, 2 * ncol(x))) + 1 ), thresh (0.001), pmethod (exhaustive), nfold (10) brt: formula (~ DD_0 + DD18 + MSP + NFFD + eFFP + PAS + RH + TD + EXT + CMD + MAR + PET), distribution (bernoulli), nTrees (10000), interactionDepth (number of predictors +1 ), shrinkage (0.01), bagFraction (0.4), trainFraction (1), cvFolds (10)

randomForest: ntree (1000), mtry (default: if (!is.null(y) && !is.factor(y)) max(floor(ncol(x)/3), 1) else floor(sqrt(ncol(x))))

<Model settings (extrapolation)>

#### Model estimates

<Coefficients>

Variable importance: We quantified the predictors' relative importance consistently across all model types. To this end, we calculated the inverse of Spearman rank correlation coefficients between probabilities of occurrence predicted by the models using permuted predictor values and predictions based on the original data. See Figure S3.7 (Supplement 3).

#### Model selection - model averaging - ensembles

Model selection: An information-theoretic approach was used for GLM, GAM, MARS (model selection based on AIC) and shrinkage for GBM.

Model averaging: Consensus predictions were generated across five model algorithms and five split-validation runs using weighted ensemble means (based on AUC).

#### Analysis and Correction of non-independence

Spatial autocorrelation: Just for glm: We did not find significant auto-correlation in residuals of models that included the selected predictor variables and their second order polynomials (but in null model). Tests were done using the R package DHARMa.

#### Threshold selection

Threshold selection: Thresholds for TSS and Kappa were optimized using the function 'Find.Optim.Stat' (R package biomod2)

#### Assessment

#### Performance statistics

Performance on training data: AIC, AUC

Performance on validation data: AUC, Kappa, TSS

<Performance on test data>

#### Plausibility check

Response shapes: Partial response curves

Expert judgement: Map display

#### Prediction

##### Prediction output

Prediction unit: Probability of occurrence, binary absence / presence prediction

Post-processing: N/A

##### Uncertainty quantification

Scenario uncertainty: For uncertainty in the climate input variables please refer to the original publications: Title, P.O. & Bemmels, J.B. (2018). ENVIREM: an expanded set of bioclimatic and topographic variables increases flexibility and improves performance of ecological niche modeling. Ecography, 41, 291–307. Wang, T., Hamann, A., Spittlehouse, D. & Carroll, C. (2016). Locally Downscaled and Spatially Customizable Climate Data for Historical and Future Periods for North America. PLOS ONE, 11, e0156720.
