## Appendix S2 for "Novel genomic offset metrics account for local adaptation in climate suitability forecasts and inform assisted migration"

Novel genomic offset metrics account for local adaptation in climate suitability forecasts and inform assisted migration

Susanne Lachmuth, Thibaut Capblancq, Anoob Prakash, Stephen R. Keller, Matthew C. Fitzpatrick

*Accessed through GBIF*

GBIF Occurrence Download https://doi.org/10.15468/dl.gtx4ii Accessed from R via rgbif (https://github.com/ropensci/rgbif) on 2020-03-30

*Accessed through BIEN*

Acadia University - ACAD. Herbarium records published by Acadia University, Canada, Nova Scotia, Wolfville. (Accessed through The Botanical Information and Ecology Network (BIEN), https://bien.nceas.ucsb.edu/bien, 2020-02-27)

Missouri Botanical Garden - MO. Herbarium records published by Missouri Botanical Garden, U.S.A., Missouri, Saint Louis. (Accessed through The Botanical Information and Ecology Network (BIEN), https://bien.nceas.ucsb.edu/bien, 2020-02-27)

The Carolina Vegetation Survey. Vegetation survey records published by The Carolina Vegetation Survey. http://cvs.bio.unc.edu/ (Accessed through The Botanical Information and Ecology Network (BIEN), https://bien.nceas.ucsb.edu/bien, 2020-02-27)

The New York Botanical Garden - NYBG. Herbarium records published by The New York Botanical Garden, U.S.A., Bronx, NY. (Accessed through The Botanical Information and Ecology Network (BIEN), https://bien.nceas.ucsb.edu/bien, 2020-02-27)

Université Laval- QFA. Herbarium records published by Université Laval, Canada, Québec, Québec. (Accessed through The Botanical Information and Ecology Network (BIEN), https://bien.nceas.ucsb.edu/bien, 2020-02-27)

VegBank: The vegetation plot archive of the Ecological Society of America. http://vegbank.org. *Published by* Peet, R.K., M.T. Lee, M.D. Jennings, D. Faber-Langendoen (eds) *(Accessed through* The Botanical Information and Ecology Network (BIEN)*, https://bien.nceas.ucsb.edu/bien, 2020-02-27)*

*Accessed through BISON*

Buffalo Society of Nature Science. *(Accessed through Biodiversity Information Serving Our Nation (BISON), https://bison.usgs.gov, 2020-02-27)*

Canadian Museum of Nature Herbarium. Canadian Museum of Nature, Canada, Ontario, Ottawa,. https://nature.ca/en/research-collections/collections. *(Accessed through Biodiversity Information Serving Our Nation (BISON), https://bison.usgs.gov, 2020-02-27)*

Consortium of California Herbaria. *(Accessed through Biodiversity Information Serving Our Nation (BISON), https://bison.usgs.gov, 2020-02-27)*

Consortium of Pacific Northwest Herbaria. *(Accessed through Biodiversity Information Serving Our Nation (BISON), https://bison.usgs.gov, 2020-02-27)*

Kathryn Kalmbach Herbarium (Denver Botanic Gardens). *(Accessed through Biodiversity Information Serving Our Nation (BISON), https://bison.usgs.gov, 2020-02-27)*

Louisiana State University Herbarium. *(Accessed through Biodiversity Information Serving Our Nation (BISON), https://bison.usgs.gov, 2020-02-27)*

Lund Botanical Museum –LD. Sweden. *(Accessed through Biodiversity Information Serving Our Nation (BISON), https://bison.usgs.gov, 2020-02-27)*

Milwaukee Public Museum. *(Accessed through Biodiversity Information Serving Our Nation (BISON), https://bison.usgs.gov, 2020-02-27)*

National Museum of Natural History, Smithsonian Institution. *(Accessed through Biodiversity Information Serving Our Nation (BISON), https://bison.usgs.gov, 2020-02-27)*

NatureServe. *(Accessed through Biodiversity Information Serving Our Nation (BISON), https://bison.usgs.gov, 2020-02-27)*

Royal Botanic Garden Edinburgh. *(Accessed through Biodiversity Information Serving Our Nation (BISON), https://bison.usgs.gov, 2020-02-27)*

Royal Botanic Gardens, Kew. *(Accessed through Biodiversity Information Serving Our Nation (BISON), https://bison.usgs.gov, 2020-02-27)*

Royal Ontario Museum. *(Accessed through Biodiversity Information Serving Our Nation (BISON), https://bison.usgs.gov, 2020-02-27)*

Staatliche Naturwissenschaftliche Sammlungen Bayerns. *(Accessed through Biodiversity Information Serving Our Nation (BISON), https://bison.usgs.gov, 2020-02-27)*

University of Connecticut. *(Accessed through Biodiversity Information Serving Our Nation (BISON), https://bison.usgs.gov, 2020-02-27)*

University of Kansas Biodiversity Institute. *(Accessed through Biodiversity Information Serving Our Nation (BISON), https://bison.usgs.gov, 2020-02-27)*

University of New Mexico Herbarium - UNM. *(Accessed through Biodiversity Information Serving Our Nation (BISON), https://bison.usgs.gov, 2020-02-27)*

Vanderbilt University. *(Accessed through Biodiversity Information Serving Our Nation (BISON), https://bison.usgs.gov, 2020-02-27)*

Yale University Peabody Museum. *(Accessed through Biodiversity Information Serving Our Nation (BISON), https://bison.usgs.gov, 2020-02-27)*

*Associated publications*

Maitner, B.S., Boyle, B., Casler, N., Condit, R., Donoghue, J., Durán, S.M., *et al.* (2018). The bien r package: A tool to access the Botanical Information and Ecology Network (BIEN) database. *Methods in Ecology and Evolution*, 9, 373–379.

Peet, R.K., Lee, M.T., Boyle, M.F., Wentworth, T.R., Schafale, M.P. & Weakley, A.S. (2012a). Vegetation-plot database of the Carolina Vegetation Survey. Biodiversity and Ecology, 4, 243–253.

Peet, R.K., Lee, M.T., Jennings, M.D. & Faber-Langendoen, D. (2012b). VegBank: a permanent, open-access archive for vegetation plot data. Biodivers. Ecol., 4, 233–241.

*R packages used for downloading occurrence data*

Chamberlain, S. (2019). rbison: Interface to the "USGS" "BISON" API. R package version 0.8.0., https://CRAN.R-project.org/package=rbison

Chamberlain, S., Barve, V., Mcglinn, D., Oldoni, D., Desmet, P., Geffert, L., et al. (2020). rgbif: Interface to the Global Biodiversity Information Facility API. R package version 1.4.0, <URL: https://CRAN.R-project.org/package=rgbif>.

Maitner, B. (2018). BIEN: Tools for Accessing the Botanical Information and Ecology Network Database. R package version 1.2.3. https://CRAN.R-project.org/package=BIEN
