## Appendix S3 for "Novel genomic offset metrics account for local adaptation in climate suitability forecasts and inform assisted migration"

Novel genomic offset metrics account for local adaptation in climate suitability forecasts and inform assisted migration

Susanne Lachmuth, Thibaut Capblancq, Anoob Prakash, Stephen R. Keller, Matthew C. Fitzpatrick

### Supplementary figures

Table of Contents:


| | |
|---|---|
| Figure S1 | Page 2 |
| Figure S2 | Page 3 |
| Figure S3 | Page 4 |
| Figure S4 | Page 5 |
| Figure S5 | Page 6 |
| Figure S6 | Page 7 |
| Figure S7 | Page 8 |
| Figure S8 | Page 9 |
| References | Page 10 |

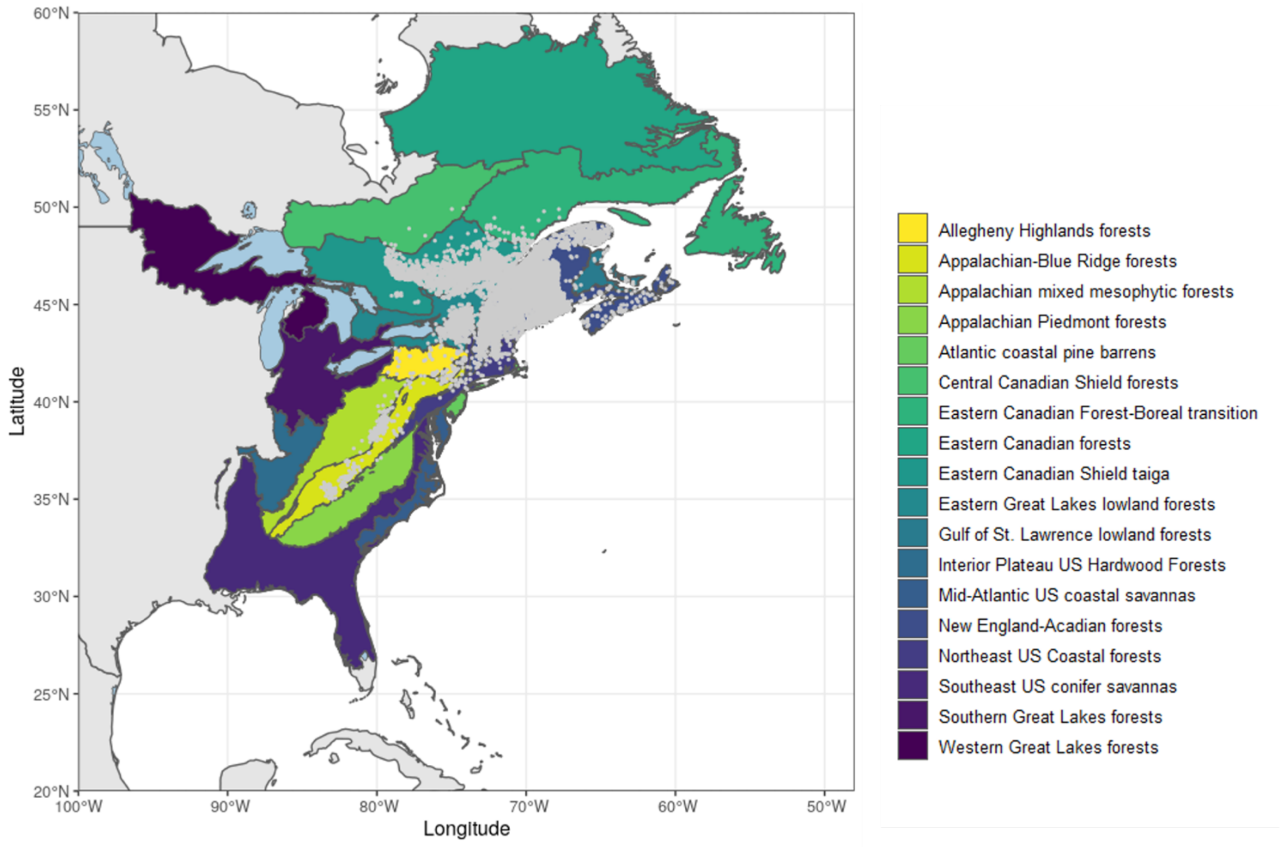


**Figure S1**: Observed presence of red spruce *Picea rubens* Sarg. in ecoregions of eastern North America at a resolution of 2.5 arc minutes. Ecoregions according to (Dinerstein et al. 2017).

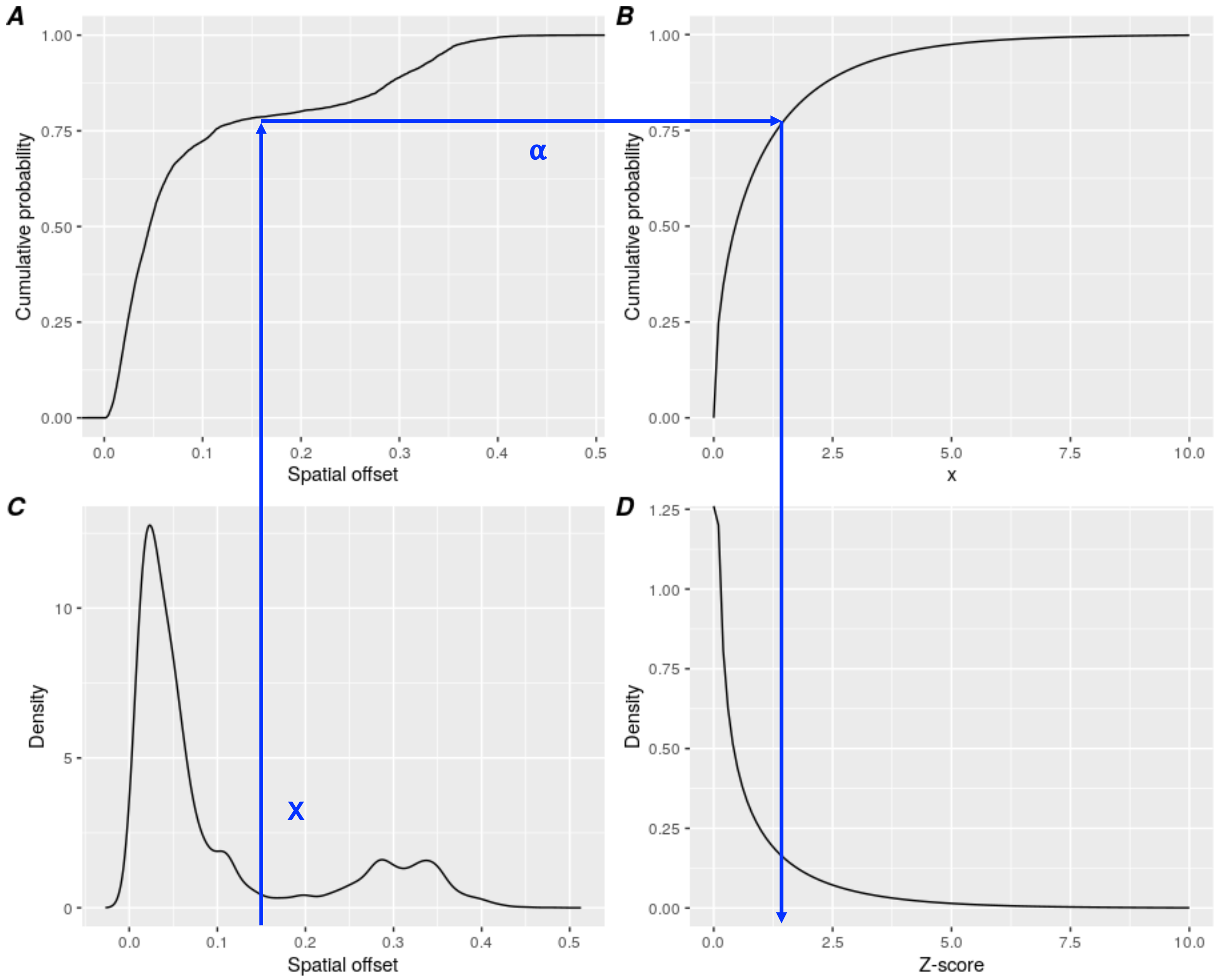


**Figure S2**: Schematic illustration of offset standardization through quantile normalization (see also Figure 1). The (spatio-) temporal offset value x (x-axis of A and C), which is the α-th quantile of an empirical distribution of contemporary spatial offsets (A), is mapped to the α quantile of the Chi distribution with one degree of freedom used here as reference distribution (C) and then re-expressed as a z-score (D, standard deviations of the reference probability distribution).

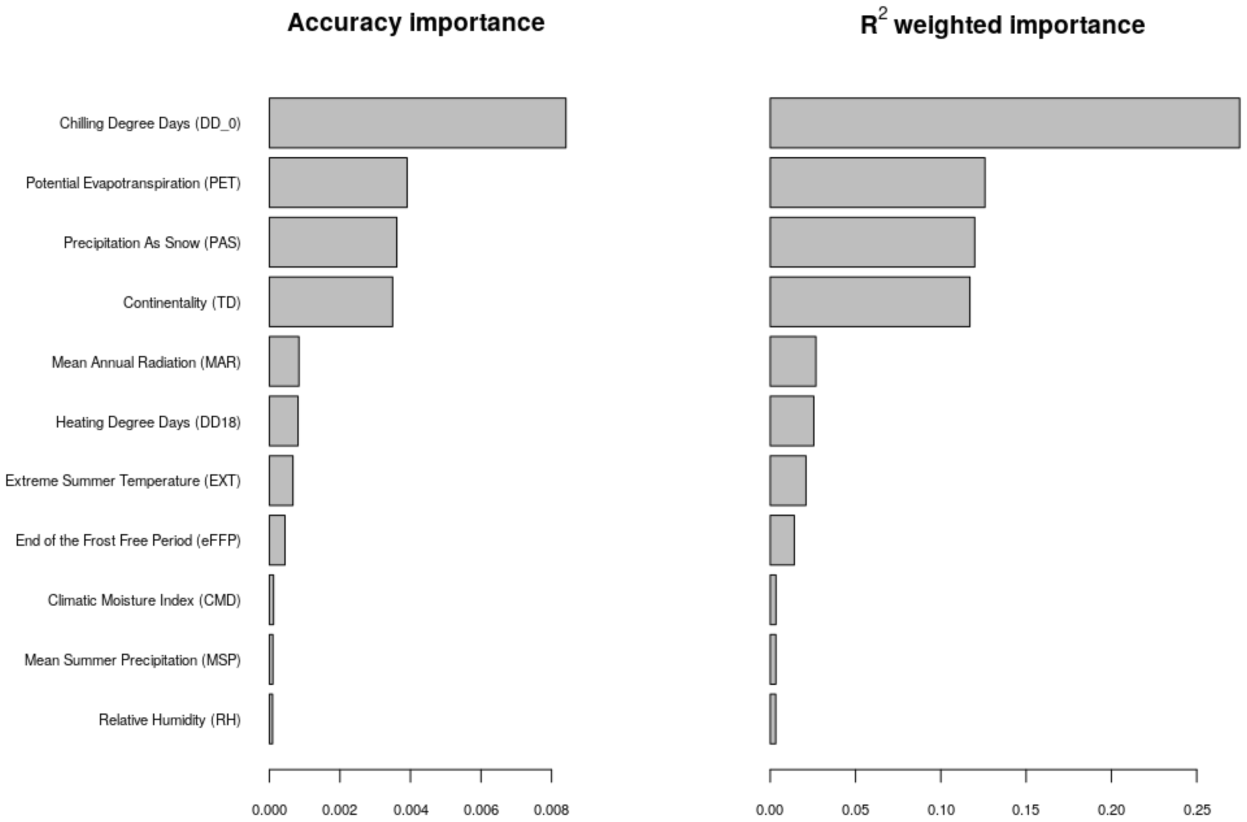


**Figure S3**: The relative importance of each climatic predictor of the Gradient Forest model in describing the genome-wide allele frequency variation along climatic gradients.

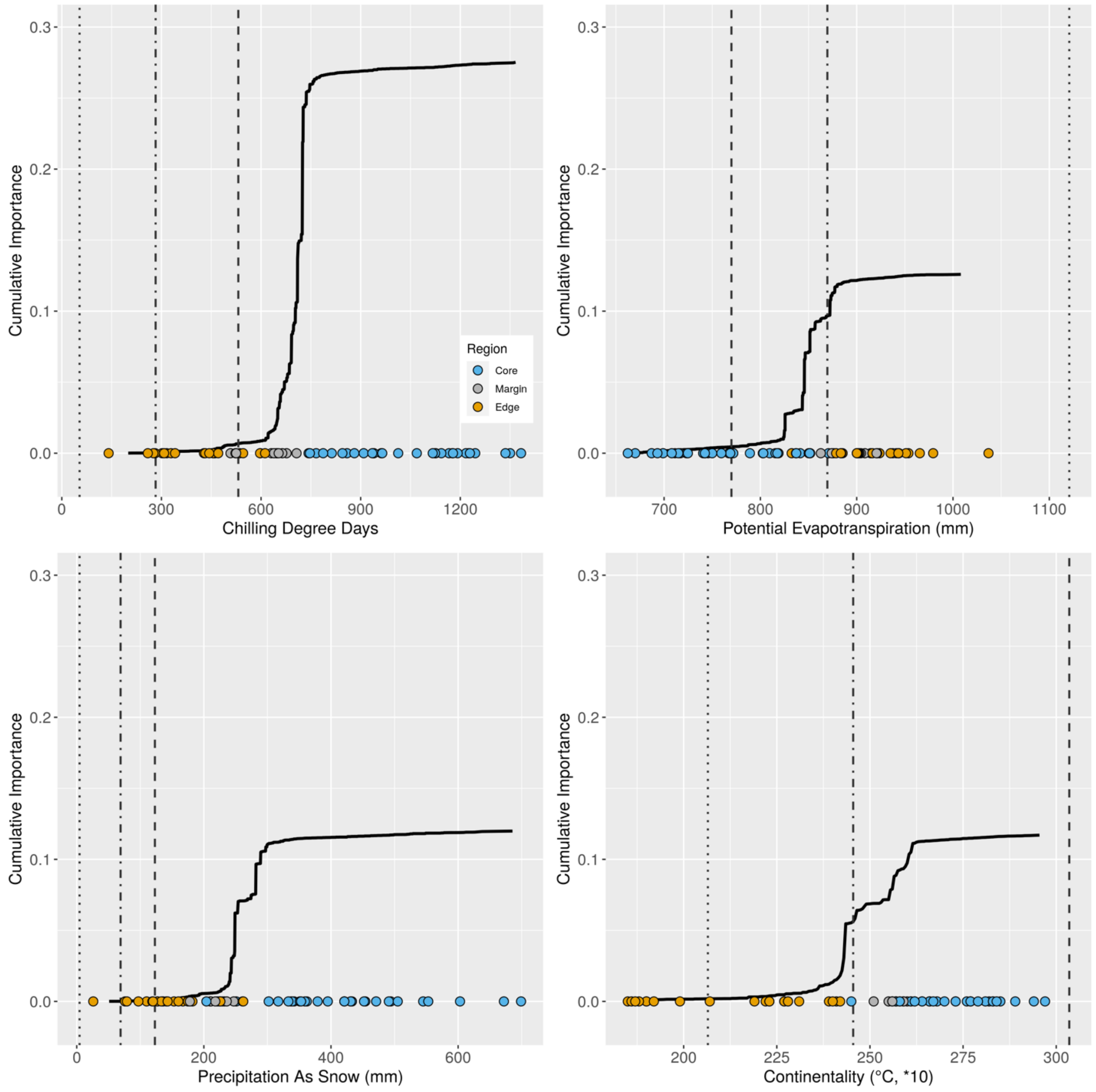


**Figure S4**: Average allele frequency turnover functions for the most important climatic predictors from the Gradient Forest model. Each bold line represents the $R^2$-weighted average of the individual turnover functions for each of the 240 putatively adaptive SNPs. Vertical lines indicate the local climatic conditions in our common gardens during the experiments in the years 2019-2020; dotted, North Carolina garden; dash-dotted, Maryland garden; dashed, Vermont garden. Dots indicate the **current climate (1961-1990 CE)** at our red spruce sampling locations in three regions.

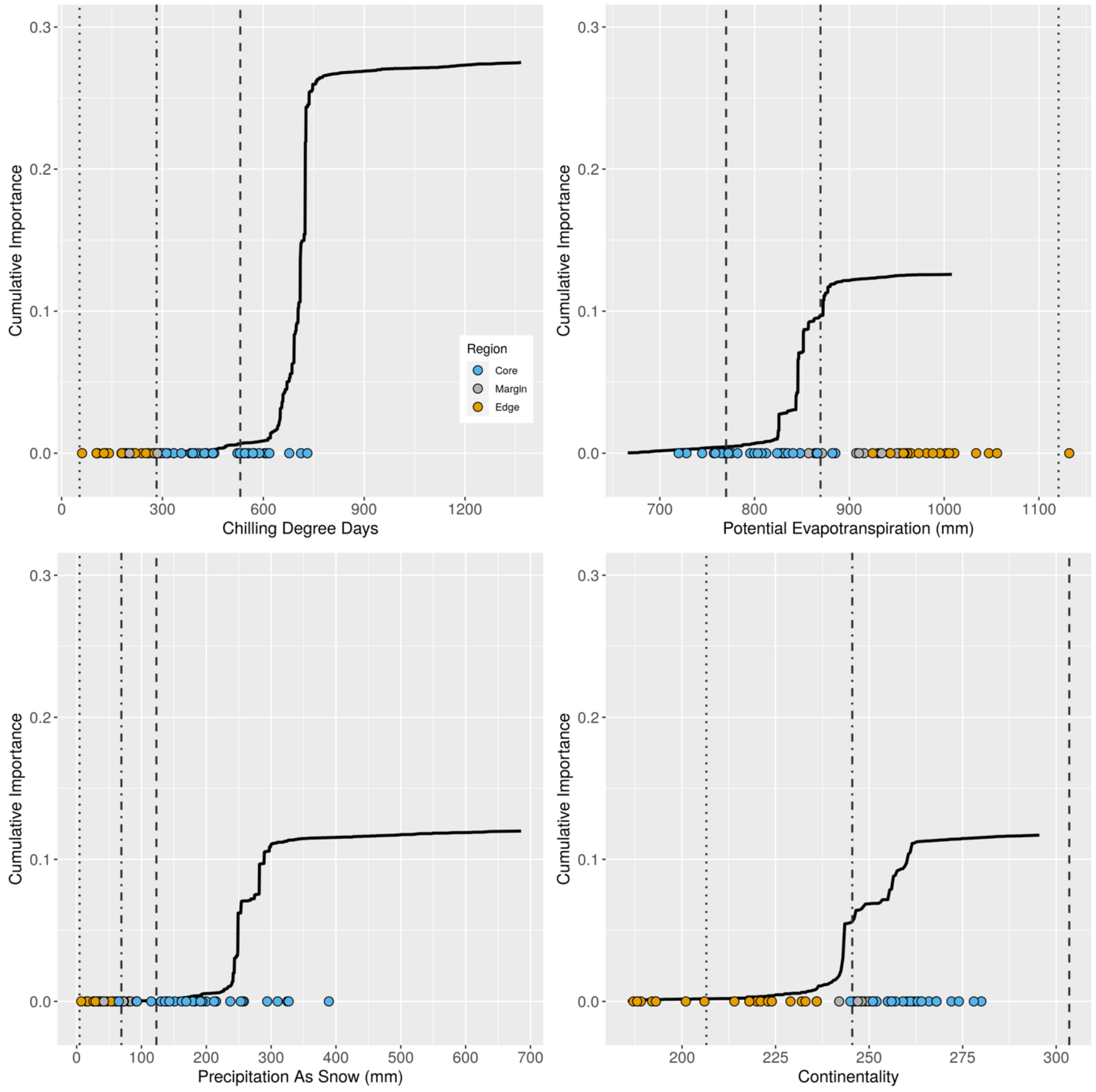


**Figure S5**: Average allele frequency turnover functions for the most important climatic predictors from the Gradient Forest model. Each bold line represents the $R^2$-weighted average of the individual turnover functions for each of the 240 putatively adaptive SNPs. Vertical lines indicate the local climatic conditions in our common gardens during the experiments in the years 2019-2020; dotted, North Carolina garden; dash-dotted, Maryland garden; dashed, Vermont garden. Dots indicate the **future climate (2071-2100 CE, under a moderate shared socio-economic pathway (SSP2-45))** at our red spruce sampling locations in three regions of its distribution range.

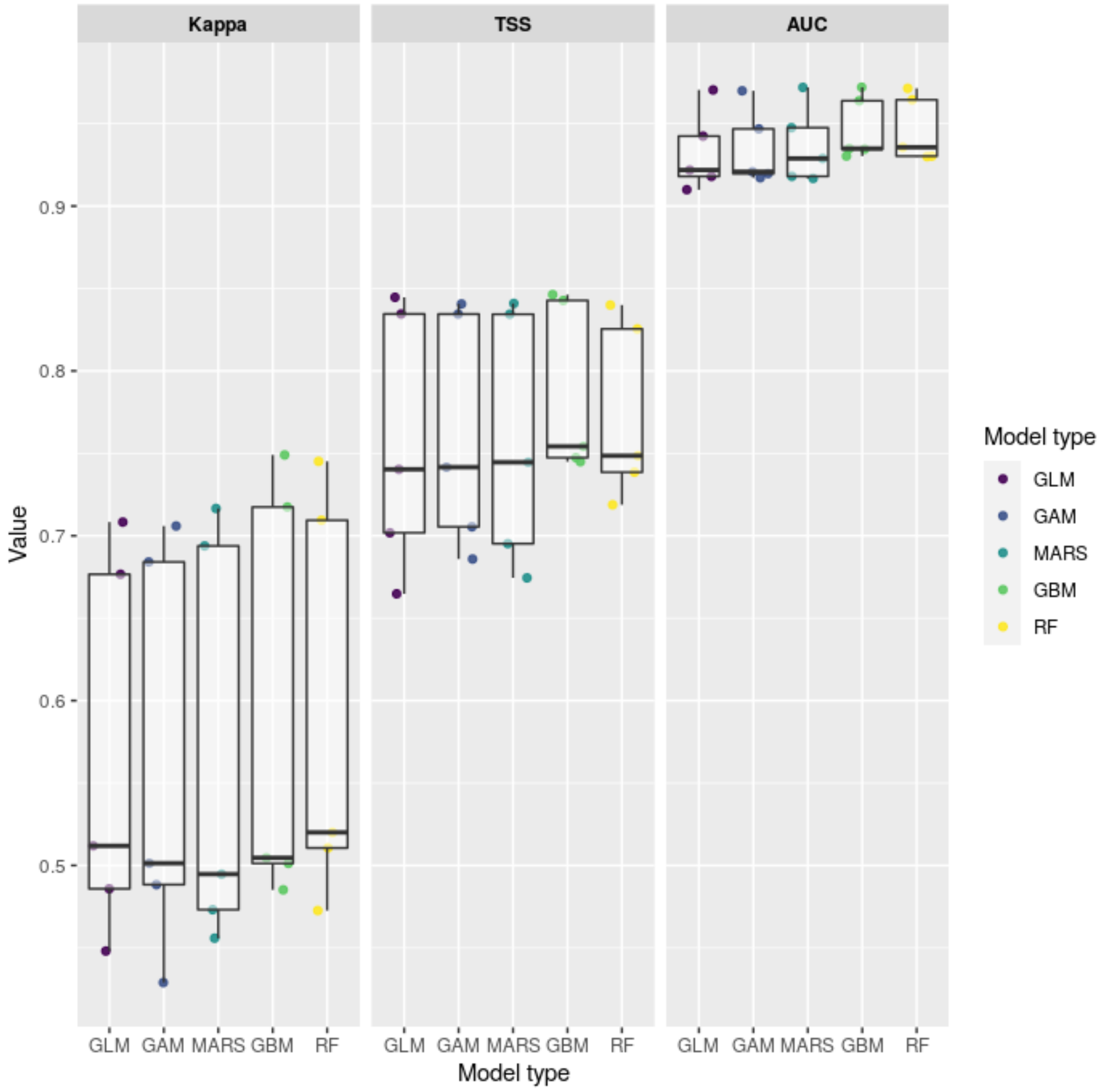


**Figure S6**: Variation in the Kappa, TSS (true skill statistics) and AUC (area under the receiver-operating characteristic curve) statistics of model accuracy across five split-validation runs for Ecological Niche Models (ENMs) using the ENM predictor set (see Table 1) and five different algorithms: generalized linear and generalized additive models (GLM, GAM), multivariate adaptive regression splines (MARS), gradient boosting machines (GBM), and random forests (RF).

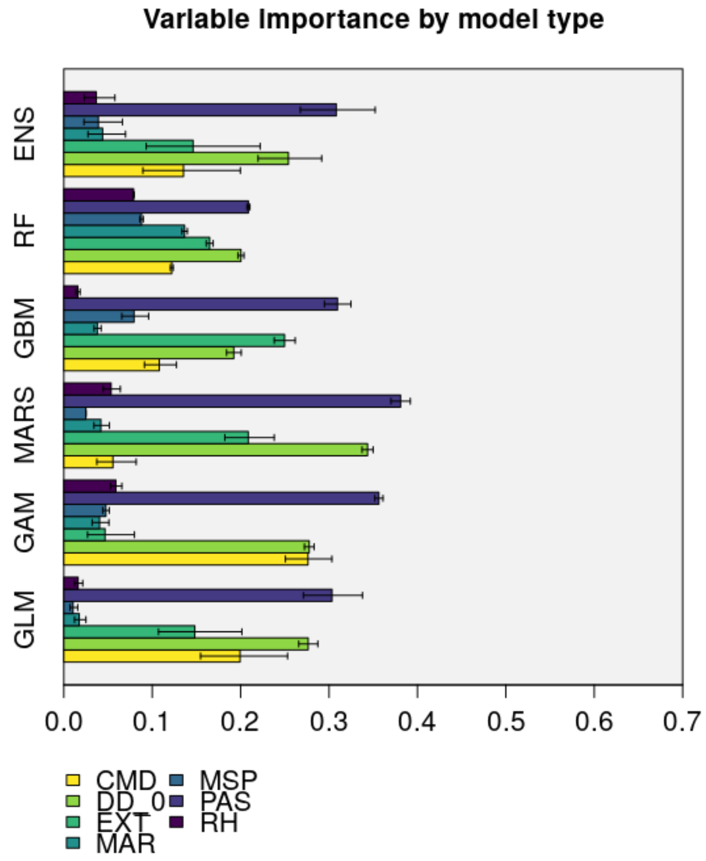


**Figure S7**: Variable importance for the Ecological Niche Model (ENM) using the ENM predictor set (see Table 1) of the ensemble and five different ecological niche modeling algorithms: generalized linear and generalized additive models (GLM, GAM), multivariate adaptive regression splines (MARS), gradient boosting machines (GBM), and random forests (RF). The bars show the mean values across five split validation runs. Error bars represent the 95% confidence intervals. CMD, Climatic Moisture Index; DD_0, Chilling degree days; EXT, extreme maximum temperature; MAR, Mean Annual Radiation; MSP, Mean Summer Precipitation; PAS, Precipitation As Snow; RH, Relative Humidity.

We quantified the predictors' relative importance consistently across all model types. To this end, we calculated the inverse of Spearman rank correlation coefficients between probabilities of occurrence predicted by the models using permuted predictor values and predictions based on the original data (Boonman et al. 2020; Thuiller et al. 2009). High values indicate high variable importance.

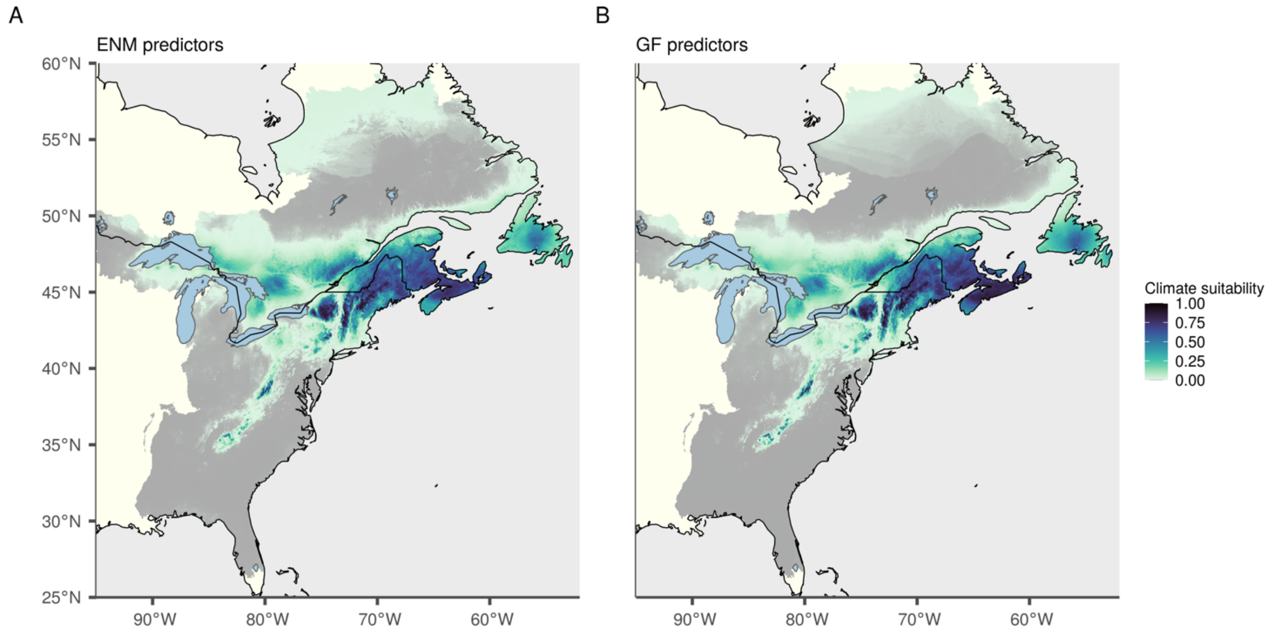


**Figure S8**: Projections of current habitat suitability (weighted ensemble mean probability of occurrence) as predicted by classic ecological niche models (ENM) based on (A) the seven climate variables of the ENM predictor set (Table 1) and (B) the eleven climate variables used also for Gradient Forest model fitting (GF predictor set) under current climate (1961-1990 CE).
