## Appendix S4 for "Novel genomic offset metrics account for local adaptation in climate suitability forecasts and inform assisted migration"

Novel genomic offset metrics account for local adaptation in climate suitability forecasts and inform assisted migration

Susanne Lachmuth, Thibaut Capblancq, Anoob Prakash, Stephen R. Keller, Matthew C. Fitzpatrick

**Results of Ecological Niche Model fitted using the Gradient Forest predictor set**

Table of Contents:


| | |
|---|---|
| Figure S1 | Page 2 |
| Figure S2 | Page 3 |
| Figure S3 | Page 4 |
| References | Page 5 |

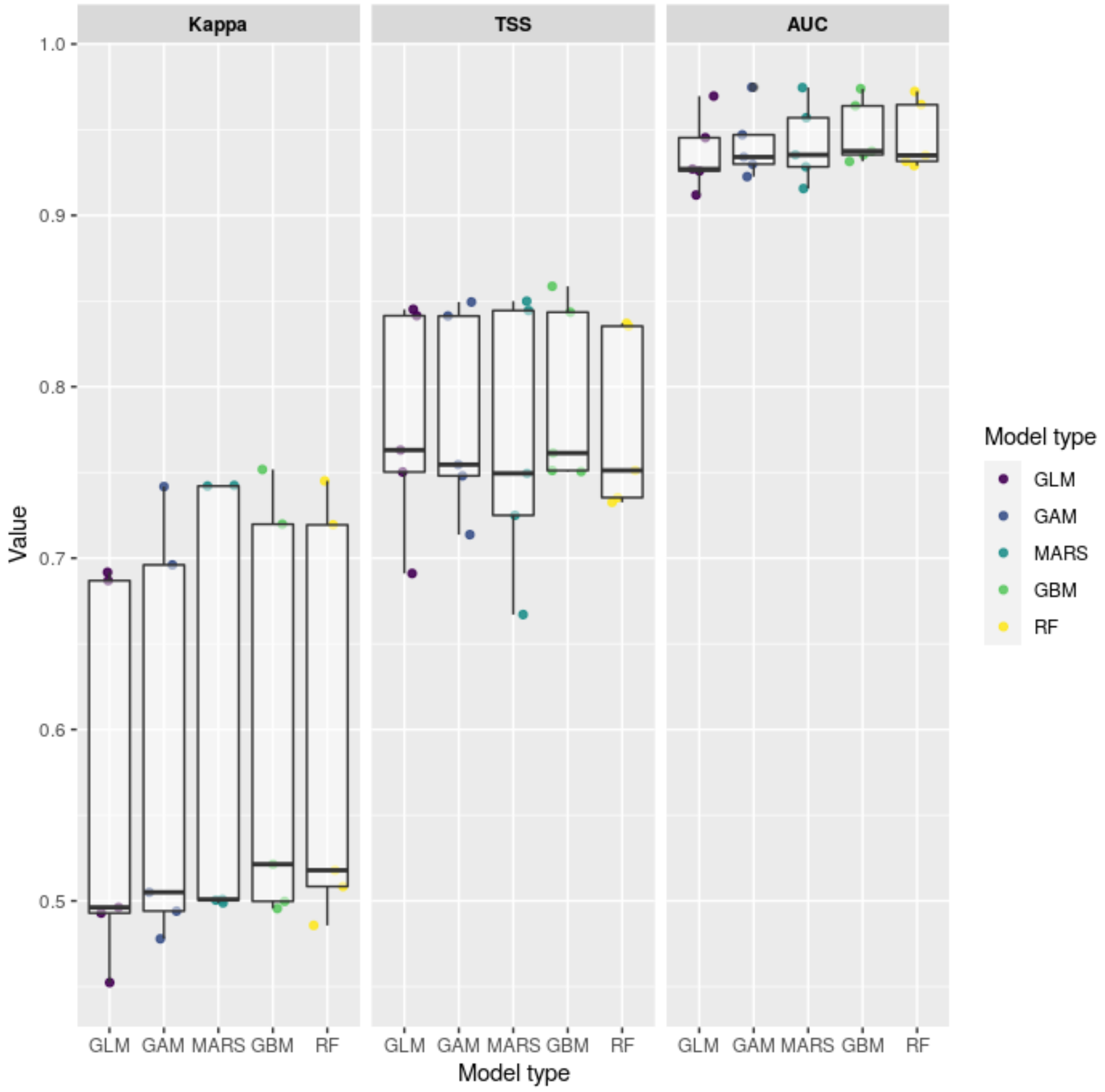


**Figure S1**: Variation in the Kappa, TSS (true skill statistics) and AUC (area under the receiver-operating characteristic curve) statistics of model accuracy across five split-validation runs for Ecological Niche Models (ENMs) using the Gradient Forest (GF) predictor set (see Table 1) and five different algorithms: generalized linear and generalized additive models (GLM, GAM), multivariate adaptive regression splines (MARS), gradient boosting machines (GBM), and random forests (RF).

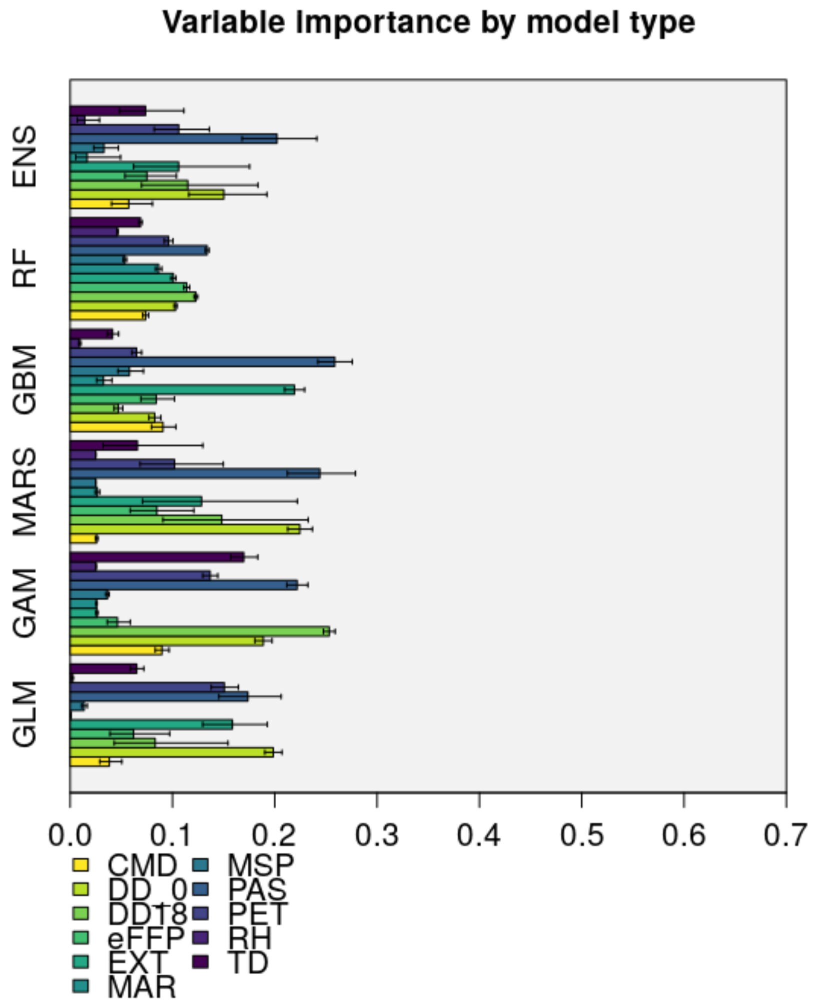


**Figure S2**: Variable importance for the Ecological Niche Model (ENM) using the Gradient Forest (GF) predictor set (see Table 1) of the ensemble and five different ecological niche modeling algorithms: generalized linear and generalized additive models (GLM, GAM), multivariate adaptive regression splines (MARS), gradient boosting machines (GBM), and random forests (RF). The bars show the mean values across five split validation runs. Error bars represent the 95% confidence intervals. CMD, Climatic Moisture Index; DD_0, Chilling degree days; DD18, Heating Degree Days; eFFP End of Frost Free Period; EXT, Extreme Maximum Temperature; MAR, Mean Annual Radiation; MSP, Mean Summer Precipitation; PAS, Precipitation As Snow; PET, Potential Evapotranspiration; RH, Relative Humidity; TD, Continentality.

We quantified the predictors' relative importance consistently across all model types. To this end, we calculated the inverse of Spearman rank correlation coefficients between probabilities of occurrence predicted by the models using permuted predictor values and predictions based on the original data (Boonman et al., 2020; Thuiller et al., 2009). High values indicate high variable importance.

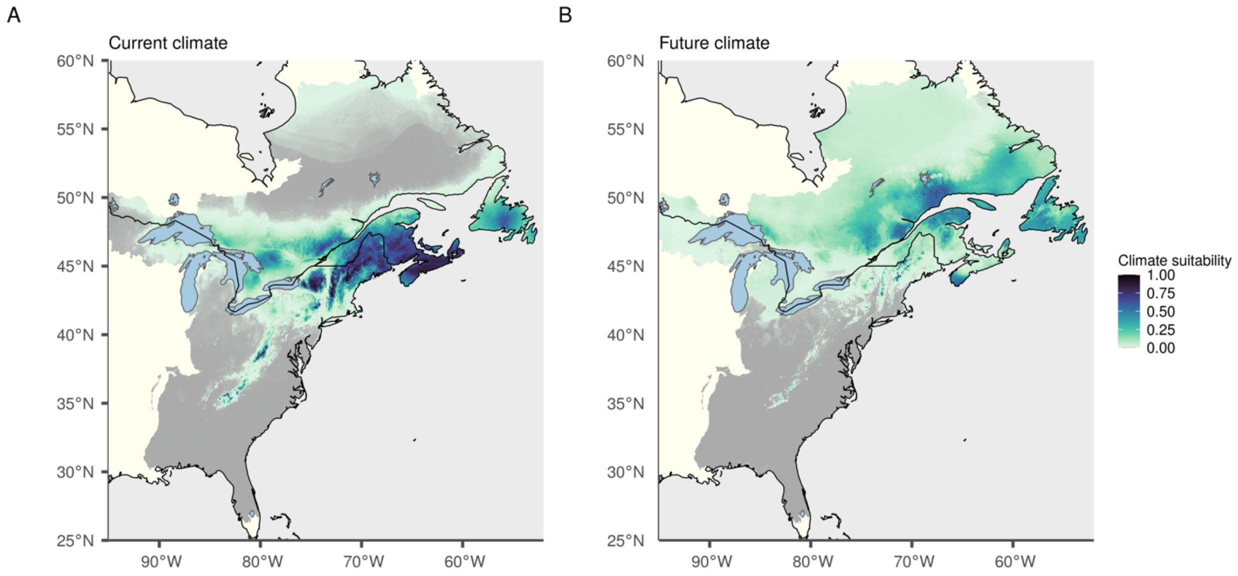


**Figure S3**: Projections of future habitat suitability for *Picea rubens* in Eastern North America for (A) current (1961-1990 CE) and (B) future climate (under the moderate shared socio-economic pathway SSP2-45, for 2071-2100 CE). The maps show weighted ensemble mean probability of occurrence as predicted by classic ecological niche models (ENM) based on the eleven climate variables used also for Gradient Forest model fitting (GF predictor set). Plot (A) is identical with Appendix S3: Figure S8 B. ENM predictions using the ENM predictor set can be found in Figure Appendix S3: Figure S8 A (current) and the main text Figure 4 A (future).
